# The evolution of mechanisms to produce phenotypic heterogeneity in microorganisms

**DOI:** 10.1101/2020.02.25.964643

**Authors:** Guy Alexander Cooper, Ming Liu, Jorge Peña, Stuart Andrew West

## Abstract

In bacteria and other microorganisms, the cells within a population often show extreme phenotypic variation. Different species use different mechanisms to determine how distinct phenotypes are allocated between individuals, including coordinated, random, and genetic determination. However, it is not clear if this diversity in mechanisms is adaptive—arising because different mechanisms are favoured in different environments—or is merely the result of non-adaptive artifacts of evolution. We use theoretical models to analyse the relative advantages of the two dominant mechanisms to divide labour between reproductives and helpers in microorganisms. We show that coordinated specialisation is more likely to evolve over random specialisation in well-mixed groups when: (i) social groups are small; (ii) helping is more “essential”; and (iii) there is a low metabolic cost to coordination. We find analogous results when we allow for spatial structure with a more detailed model of cellular filaments. More generally, this work shows how diversity in the mechanisms to produce phenotypic heterogeneity could have arisen as adaptations to different environments.

## INTRODUCTION

Different species use different mechanisms to produce adaptive phenotypic heterogeneity (Fig. 1)^1–5^. In some cases, there is coordination across individuals to determine which individual will perform which role (*coordinated specialisation*)^1,6^. This coordination could use signals, cues, or a developmental programme to provide information about the phenotypes adopted by other individuals in the group^7^. For example, when honey bee workers feed royal jelly to larvae to produce reproductive queens (Fig. 1A), or when the local density of a signalling molecule determines whether cyanobacteria cells develop into sterile nitrogen-fixing heterocysts (Fig. 1B)^8–10^. In other cases, each individual adopts a helper phenotype with a certain probability, independently and without knowledge of the phenotypes adopted by other individuals (*random specialisation*)^2,5,11,12^. For example, in *Salmonella enterica* co-infections, random biochemical fluctuations within each cell’s cytoplasm are used to determine whether the cell sacrifices itself to trigger an inflammatory response that eliminates competitor species (Fig. 1D)^12,13^. In yet further cases, phenotype is influenced by the individual’s genotype (genetic control). For instance, in some ant societies, whether individuals develop into queens, major or minor workers can be determined, in part, by their genes (Fig. 1C)^3,14–16^. Across the tree of life some species employ one mechanism to produce phenotypic heterogeneity whereas in other species mixed forms exist with a combination of coordinated specialisation, random specialisation, or genetic control^3,15,17–22^.

**Figure 1.**
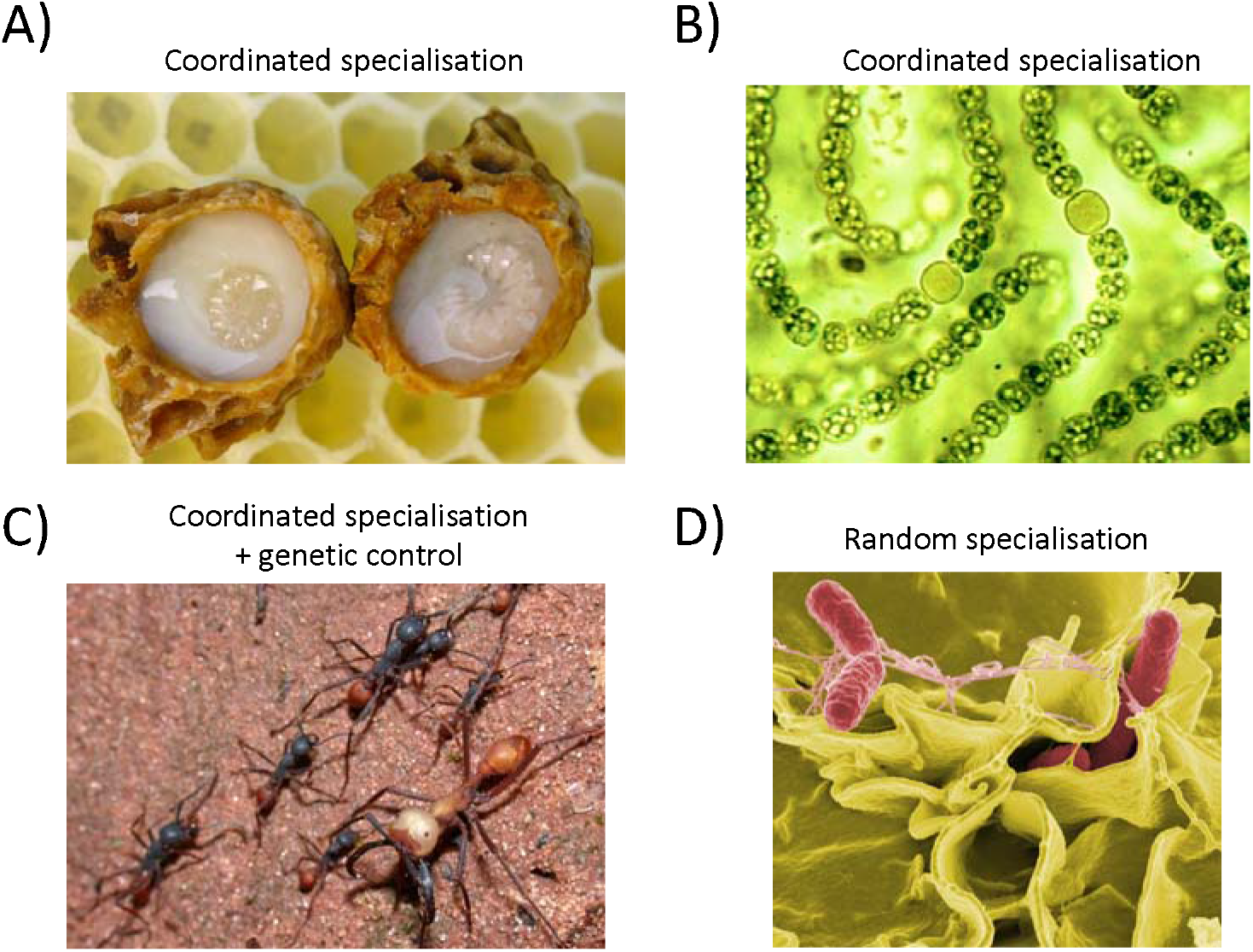
Different mechanisms to produce phenotypic heterogeneity in nature. A) In honey bee hives (A*pis mellifera*), larvae develop as sterile workers unless they are fed large amounts of royal jelly by adult workers (coordinated specialisation)^8^ (Photo by Wausberg via the Wikimedia Commons.) B) In *A. cylindrica* filaments (cyanobacteria), some individuals develop into sterile nitrogen fixers (larger, round cells) if the amount of nitrogen fixed by their neighbours is insufficient (coordinated specialisation). This leads to a precise allocation of labour, with nitrogen-fixing cells distributed at fixed intervals along the filament^9^ (Picture taken by Robert Calentine.) C) In the army ant (*Eciton Burchelli*), whether individual ants become a major or minor worker has a genetic component (genetic control)^16^ (Photo by Alex Wild via the Wikimedia Commons, cropped.) D) In *S. enterica* infections (serovar Typhymurium), each cell amplifies intra-cellular noise to determine whether it will self-sacrifice and trigger an inflammatory response that eliminates competing strains (random specialisation)^13^ (Photo by Rocky Mountain Laboratories, NIAID, NIH via Wikimedia Commons.)

We lack general evolutionary explanations for why different species use different mechanisms to produce phenotypic heterogeneity^2,3,23,24^. Previous work has focused on non-reproductive division of labour in the social insects, and the proximate mechanisms that lead to different worker castes^6,16,25–29^. However, the focus in that literature is on a different question – how different proximate mechanisms can produce coordinated specialisation – rather than the broader question of whether coordinated specialisation should be favoured over random specialisation or genetic control in the first place. It is with reproductive division of labour that these three very different mechanisms have been observed in different species and for which there is an absence of evolutionary explanations^2,3,23,24,30^.

Reproductive division of labour in bacteria and other social microbes offers an excellent opportunity for studying why different mechanisms to produce phenotypic heterogeneity are favoured in different species^1,2^. Reproductive division of labour occurs when social groups are composed of more cooperative ‘helpers’ who gain indirect fitness benefits by the aid they provide to less cooperative ‘reproductives’. Across microbes, the two primary mechanisms used to produce reproductive division of labour are coordinated and random specialisation (Fig. 2). Furthermore, while the form of cooperation and life histories of social microbes share many similarities, they also vary in factors that could influence the evolution of division of labour, such as social group size ^31,32^.

**Figure 2.**
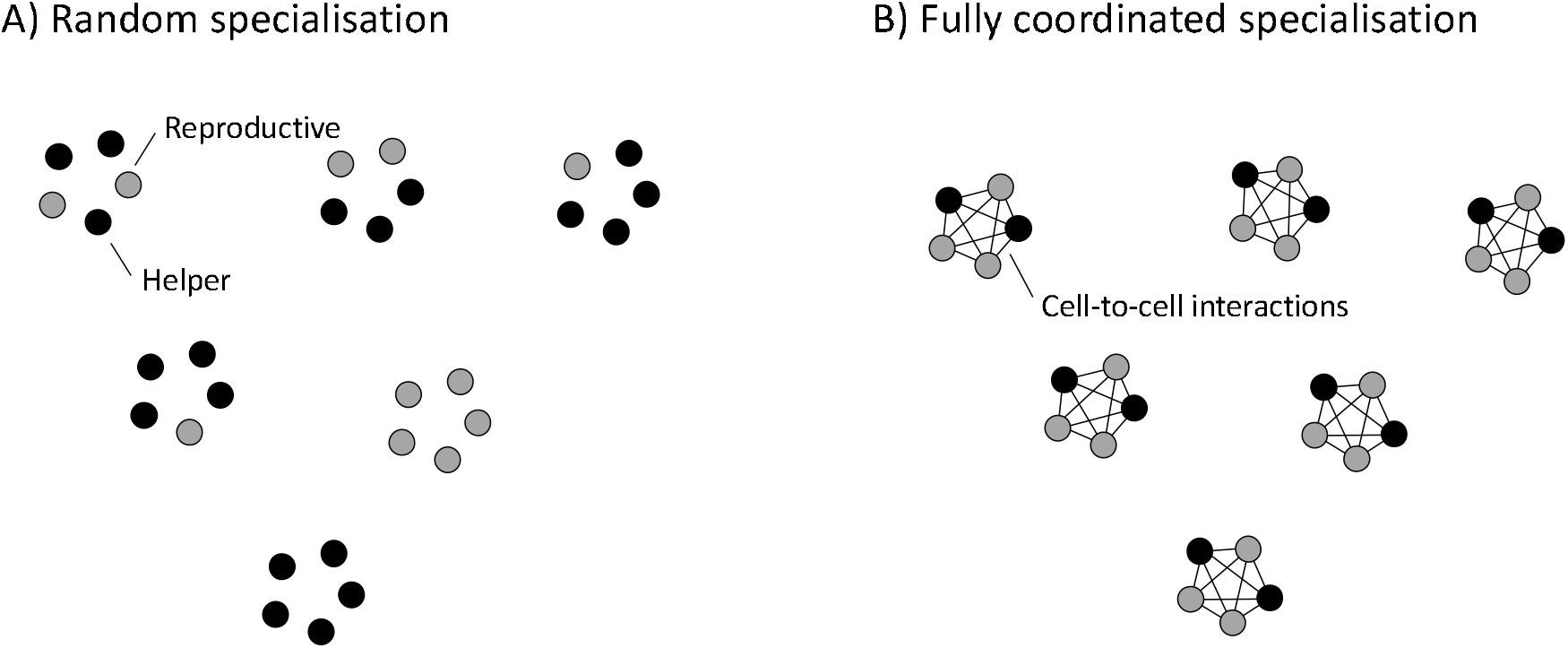
Mechanisms to produce reproductive division of labour in clonal groups. We examine the relative advantages and disadvantages of the two key mechanisms to produce reproductive division of labour in social microorganisms^1,5,11,33^. (A) Random specialisation occurs when cells randomly specialise into helpers or reproductives independently of one another. This can occur when a genetic feedback circuit is used to amplify small molecular fluctuations in the cytoplasm of each cell (phenotypic noise) ^4,11,12,34–36^. (B) Coordinated specialisation occurs when cells interact with one another, and share (or gain) phenotypic information while they are differentiating. This could occur through the secretion and detection of extracellular molecules (signals or cues), or with a shared developmental programme (epigenetics)^1,2,25^.

We develop theoretical models to examine whether the relative advantages of random and coordinated specialisation can depend upon social or environmental conditions. Our aim here is to use reproductive division of labour in microbes as a ‘test system’ to address the broader question of whether evolutionary models can explain the diversity in the mechanisms that produce phenotypic heterogeneity more broadly.

## RESULTS

We compare the relative fitness advantages of reproductive division of labour with either coordinated or random specialisation. Our first aim is to capture the problem in a deliberately simple model, which is easy to interpret, and can be applied across diverse microbe species^37,38^.

We begin by assuming that coordinated specialisation always produces the optimal proportion of helpers and reproductives (fully coordinated specialisation) and that there is no within-group spatial structure (well-mixed groups). We then test the robustness of our results by examining several alternate models for different biological scenarios and by developing a more detailed model of growing cyanobacteria filaments that includes the effects of within-group spatial structure. Throughout, we assume a form of cooperation that is common in microbes, where some individuals produce a ‘public good’ that benefits all cells.

### Random specialisation vs fully coordinated specialisation

We assume that a single cell arrives on an empty patch and, through a fixed series of replications, produces a clonal group of n individuals that consists of *k* sterile helpers and *n* − *k* pure reproductives (*k* ∈ {0,1,2, …, *n*). We denote group fecundity, *g*_*k,n*_, as the reproductive success of a particular group in the absence of mechanism costs. This is measured as the per capita number of offspring that would disperse at the end of the group life cycle, given by

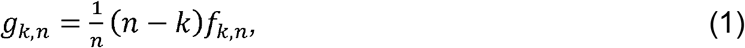

where *n* − *k* is the number of reproductives in the group, and *f*_*k,n*_ is the fecundity of each reproductive in the absence of mechanism costs. We assume that *f*_*k,n*_ increases with the number of helpers in the whole group (*k*).

Expression (1) highlights the trade-off between the number of reproductives in the group (*n* − *k*), which is higher when there are fewer helpers (lower *k*), and the amount of help that those reproductives obtain (*f*_*k,n*_), which is higher when there are more helpers (higher *k*). The balance of this trade-off often results in an optimal number of helpers, *k*\*, that is intermediate (i.e., 0 < k* < *n*), giving *g*_*k*\*,*n*_ as the maximal reproductive success of the group.

In species that divide labour by coordination, the outcome of individual specialisation depends on the phenotypes of social group neighbours. Our first model is deliberately agnostic to the details of how phenotype information is shared between group members in order to facilitate predictions across different systems. For instance, individuals may share phenotype information via signalling between cells or with a common developmental programme (Fig. 2B)^1,2,39^. We make the simplifying assumption that individuals coordinate fully, so that coordinated groups always form with precisely the optimal number of helpers, *k*\*. The disadvantage of coordinated specialisation is that the mechanism could incur metabolic costs, such as the production of extracellular signalling molecules. The fitness of a group of coordinated specialisers is given by:

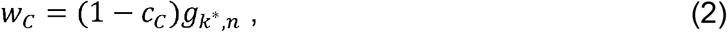

where *g*_*k*\*,*n*_ is the group fecundity with the optimal number of helpers, *k*\*, and 0≤ *C*_*c*_ ≤1 is the metabolic cost of coordination, whose form we leave unspecified but could in principle depend on further factors such as group size. A number of different models have examined how different proximate mechanisms can produce coordinated division of labour in specific systems ^6,25,28,29^.

In species that divide labour by random specialisation, each individual in the group independently becomes a helper with a given probability and a reproductive otherwise (Fig. 2A). Hence, the final number of helpers in the group is a binomial random variable. We assume that the probability of adopting a helper role is equal to the optimal proportion of helpers (*P*\* = *k*\*/*n*). Thus, the expected fitness of a group of random specialisers is given by:

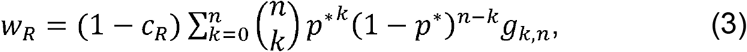

where 0≤ *c*_*R*_ ≤1 is the metabolic cost of random specialisation, which we assume is independent of the number of helpers in the group, *k*. The potential advantage of random specialisation is that there may be fewer upfront metabolic costs from, for example, between cell signalling (i.e., if *c*_*R*_ < *c*_*C*_ holds). The downside of random specialisation is that groups form most of the time with fewer or more helpers than is optimal (developmental stochasticity). In principle, the probability of becoming a helper could be transiently regulated by environmental cues to produce on average more or fewer helpers when this is more favourable. However, throughout our analysis we assume a stable environment and ignore such regulation.

We need to specify how reproductive fecundity depends on the number of helpers in the group. We focus here on one of the most common forms of cooperation in microbes, where individuals secrete factors that provide a benefit to the local population of cells (“public goods”) ^40^. We assume that the amount of public good in the social group depends linearly on the number of helpers in the group and is “consumed” by all group members equally ^41,42^. An example of such a public good is found in *Bacillus subtilis* populations, where only a subset of cells (helpers) produce and secrete proteases that degrade proteins into smaller peptides, but where these are then re-absorbed as a nutrient source by all cells ^43^.

We allow the relative importance of producing public goods to vary between species. Each reproductive has a baseline fecundity, *b* ≥ 0, that is independent of the amount of public good in the group. The fecundity benefit of helpers scales according to *h* ≥ 0 as the amount of public good in the group increases. When reproductives have no baseline fecundity (*b*= 0), we say that cooperation is essential. When baseline fecundity is non-zero (*b*> 0), cooperation is non-essential and the ratio *h*/*b* provides a useful metric for the relative importance of cooperation.

Our assumptions give the following expression for the fecundity of a reproductive:

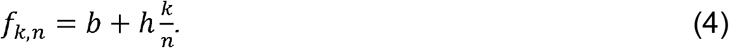

By substituting Equation 4 into Equations 1–3, we can determine when the fitness of coordinated specialisation is greater than the fitness of random specialisation (i.e., *w*_*C*_ > *w*_*R*_), which gives the simplified condition:

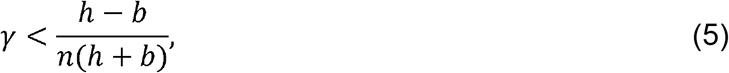

where *γ* = (*c*_*C*_ − *c*_*R*_)/(1 − *c*_*R*_) captures the relative change in metabolic costs paid when switching to coordinated specialisation from random specialisation. If *h* < *b*, then sterile helpers are disadvantageous and the group is composed of all reproductives (*k*\* = 0). Thus, division of labour with sterile helpers is favoured to evolve only when *h* > *b*, which we will assume henceforth (Supplementary Section C). Condition (5) specifies that coordinated specialisation is favoured when the relative change in metabolic costs of switching from random specialisation to coordination (*γ*), is less than the fecundity benefits gained from doing so (right-hand side). The condition can be used to predict how key environmental and ecological factors will influence which labour-dividing mechanism is more likely to evolve (Fig. 3).

**Figure 3.**
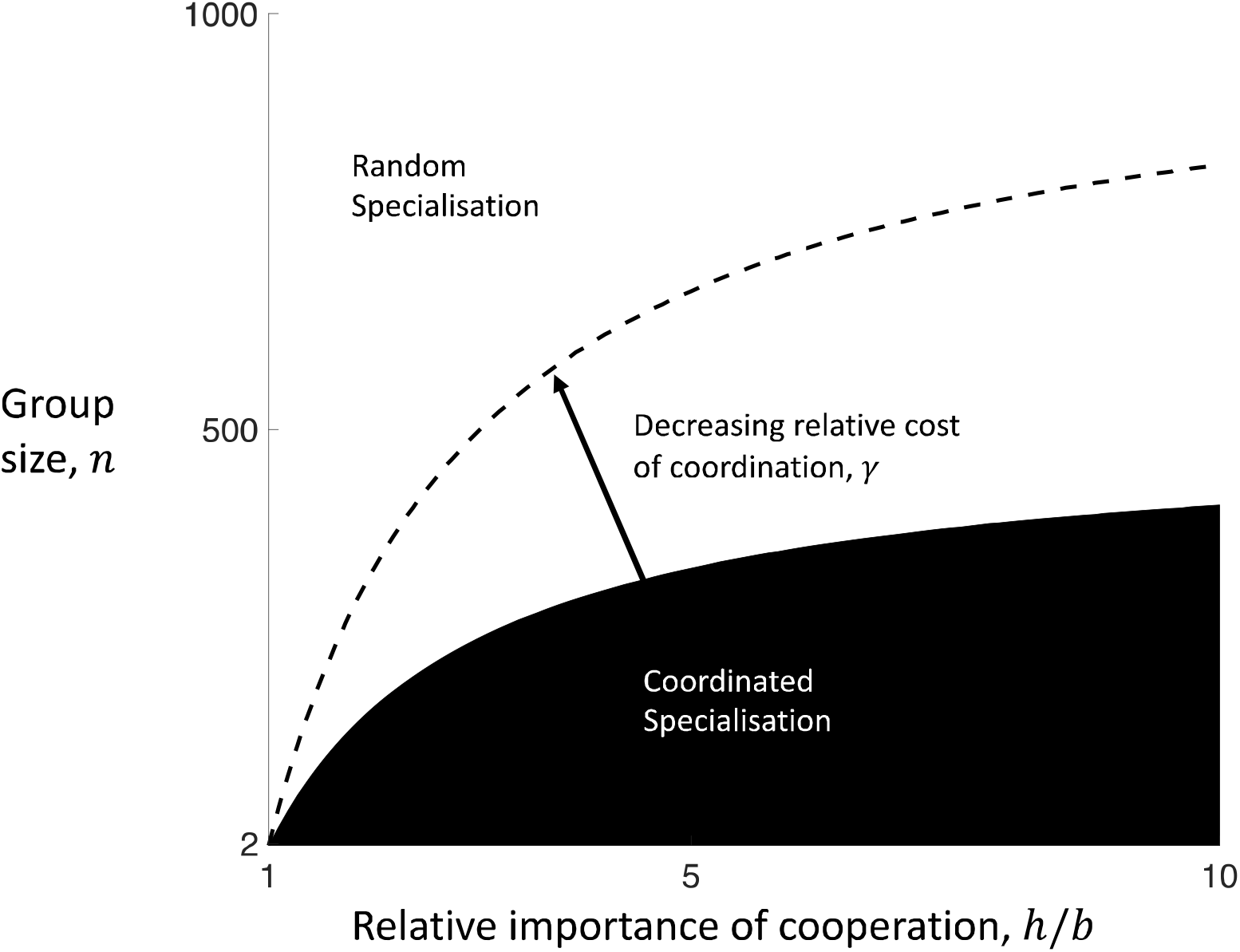
Random versus coordinated specialisation. Small group sizes (lower *n*), relatively more important cooperation (higher *h*/*b*), and lower relative metabolic costs to coordination (lower *γ*) favour division of labour by coordinated specialisation (black) over division of labour by random specialisation (white). Here we have used *γ* = 2 × 10^−3^ (solid boundary) and *γ* = 1 × 10^−3^ (dashed boundary). We note that the limit as the relative importance of cooperation goes to infinity (very large *h*/*b*) converges to the outcome for when cooperation is essential (*b*= 0).

#### Prediction 1: Smaller relative metabolic costs of coordination favour coordinated specialisation

When the metabolic cost of coordination is smaller (lower *c*_*C*_) and the metabolic cost of random specialisation is larger (higher *c*_*R*_), then the relative cost of switching from random specialisation to coordinated specialisation is lessened (smaller *γ*), which favours the evolution of coordinated specialisation (smaller left-hand side of Condition 5). If the metabolic costs of random specialisation are equal to or larger than the metabolic costs of coordination (*c*_*R*_ ≥ *c*_*C*_ ⇒ *γ* ≤ 0), then coordinated specialisation is always the favoured mechanism (Condition 5 always satisfied). Conversely, random specialisation can only ever be the favoured strategy ( *w*_*R*_ > *w*_*C*_; Condition 5 not satisfied) if the metabolic costs of random specialisation are less than the metabolic costs of coordination (*c*_*C*_ > *c*_*R*_ ⇒ *γ* > 0; a necessary but not sufficient condition). This arises directly from our starting assumption that coordinated specialisation always produces groups with the optimal proportion of helpers whereas random specialisation may often produce groups that are sub-optimal.

Larger metabolic costs of coordinated specialisation ( *c*_*R*_ < *c*_*C*_ ⇒ *γ* > 0) may be a reasonable assumption for many biological systems. The metabolic costs of random specialisation are determined by the production costs of the regulatory proteins employed in the genetic feedback circuit that amplifies intra-cellular noise^4,5,33,44^. In contrast, coordinated specialisation requires both an intracellular genetic feedback circuit and some mechanism by which phenotype is communicated between cells, such as the costly production and secretion of extra-cellular signalling molecules^1,2,9,39,45,46^.

As a result, in many scenarios, the optimal mechanism to divide labour depends on how the potentially higher metabolic costs of coordination (*c*_*C*_ > *c*_*R*_ ⇒ *γ* > 0) balance against the benefit of avoiding the stochastic costs of random specialisation (right-hand side of Condition 5). The stochastic costs of random specialisation are determined entirely by: (i) the relative likelihood that random groups deviate from the optimal proportion of helpers, and (ii) the degree to which those deviations from the optimal proportion of helpers leads to a reduced fecundity for the group (Supplementary Section C). Equation (5) shows how the importance of these two factors depends upon the size of the group (*n*) and on the relative importance of cooperation (*h*/*b*).

#### Prediction 2. Smaller social groups favour coordinated specialisation

The number of cells in the group has a large impact on the relative likelihood that random groups deviate from the optimal proportion of helpers (Fig. 3). In smaller groups, there are fewer possible outcomes for the proportion of helpers (*n*− 1 possible allocations of labour for groups of size *n*). Consequently, random specialisation can more easily lead to the formation of groups with a realised proportion of helpers that deviates significantly from the optimum (*p* ≪ *p*\* ***or*** *p* ≫ *p*\*). In contrast, in larger groups there are more possible outcomes and the resulting proportion of helpers will be more closely clustered about the optimal composition with highest fitness (with *p* ≈ *p*\* for very large group sizes).

This effect of group size on the stochastic cost of random specialisation is a consequence of the law of large numbers. For example, outcomes close to 50% heads are much more likely when tossing 100 coins in a row compared to only tossing 4 coins in a row where no heads or all heads may frequently occur. Our prediction is related to how, when mating occurs in small groups, small brood sizes select for more precise and less female biased sex ratios as there would otherwise be a high probability of producing a group containing no males at all^47–49^. In another analogue, Wahl showed a mechanistically different effect when division of labour is determined genetically and the number of group founders is small: groups may sometimes form that do not contain all of the genotypes required to produce all of the necessary phenotypes in the division of labour ^24^.

#### Prediction 3: The higher the relative importance of cooperation, the more coordinated specialisation is favoured

When the relative importance of cooperation is larger (higher *h*/*b*), the fitness costs incurred from producing too few helpers increases. In addition, as the relative importance of cooperation increases (higher *h*/*b*), the optimal proportion of helpers increases to 50% helpers 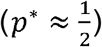. This increases the variance in the proportion of helpers produced by random specialisers, and so sub-optimal groups may arise even more frequently (Supplementary Section C). Thus, a higher relative importance of cooperation increases both (i) the likelihood that groups deviate from the optimal proportion of helpers and (ii) the scale of the fitness cost when they do. Both of these effects increase the stochastic costs of random specialisation (larger right-hand side of Condition 5), and thus favour the evolution of coordinated specialisation (Fig. 3).

### Alternative forms of cooperation

The above analysis employs a deliberately simple public goods model, focusing on factors that are expected to be relevant across many microbial systems. This facilitates the interpretation of our results and generates broadly applicable predictions that are less reliant on the details of particular species.

In order to test the robustness of our results (predictions 1–3) we also developed a series of alternative simplified models corresponding to different biological scenarios (Supplementary sections D and E; Supplementary Figs. 1-3). We examined the possibility that the public good provided by helpers: (i) is not consumed by its beneficiaries, as may occur when self-sacrificing *S. enterica* cells enter the gut to trigger an immune response that eliminates competitors (non-rivalrous or non-congestible collective good); or (ii) is only consumed by the reproductives in the group, as may preferentially occur for the fixed nitrogen secreted by heterocyst cells in *A. cylindrica* filaments (excludible or club good) ^9,12,50,51^. We allowed for reproductive fecundity to depend non-linearly on the proportion of helpers in the group, for helpers to have some fecundity (non-sterile helpers), and for division of labour to occur in each generation of group growth. In all of these alternative scenarios, we found broad qualitative agreement across the three predictions of the linear public goods model.

We found that less specialised helpers (with some fecundity) favour random specialisation over coordinated specialisation. In contrast to prediction 3, more fecund helpers can lead to a scenario where a larger relative importance of cooperation (higher *h*/*b*) disfavours coordinated specialisation. This occurs because a high relative importance of cooperation (higher *h*/*b*) can produce groups composed predominantly of non-sterile helpers (*p*\* ≈ 1), where the likelihood that random groups deviate from the optimal proportion of helpers is significantly diminished (Supplementary Section E.4).

In Supplementary Section F we develop an individual based simulation, which also supports predictions 1–3. In addition, this simulation shows that costly coordination can evolve incrementally from random specialisation, and that intermediate levels of coordination can be favoured (Supplementary Fig. 4).

**Figure 4:**
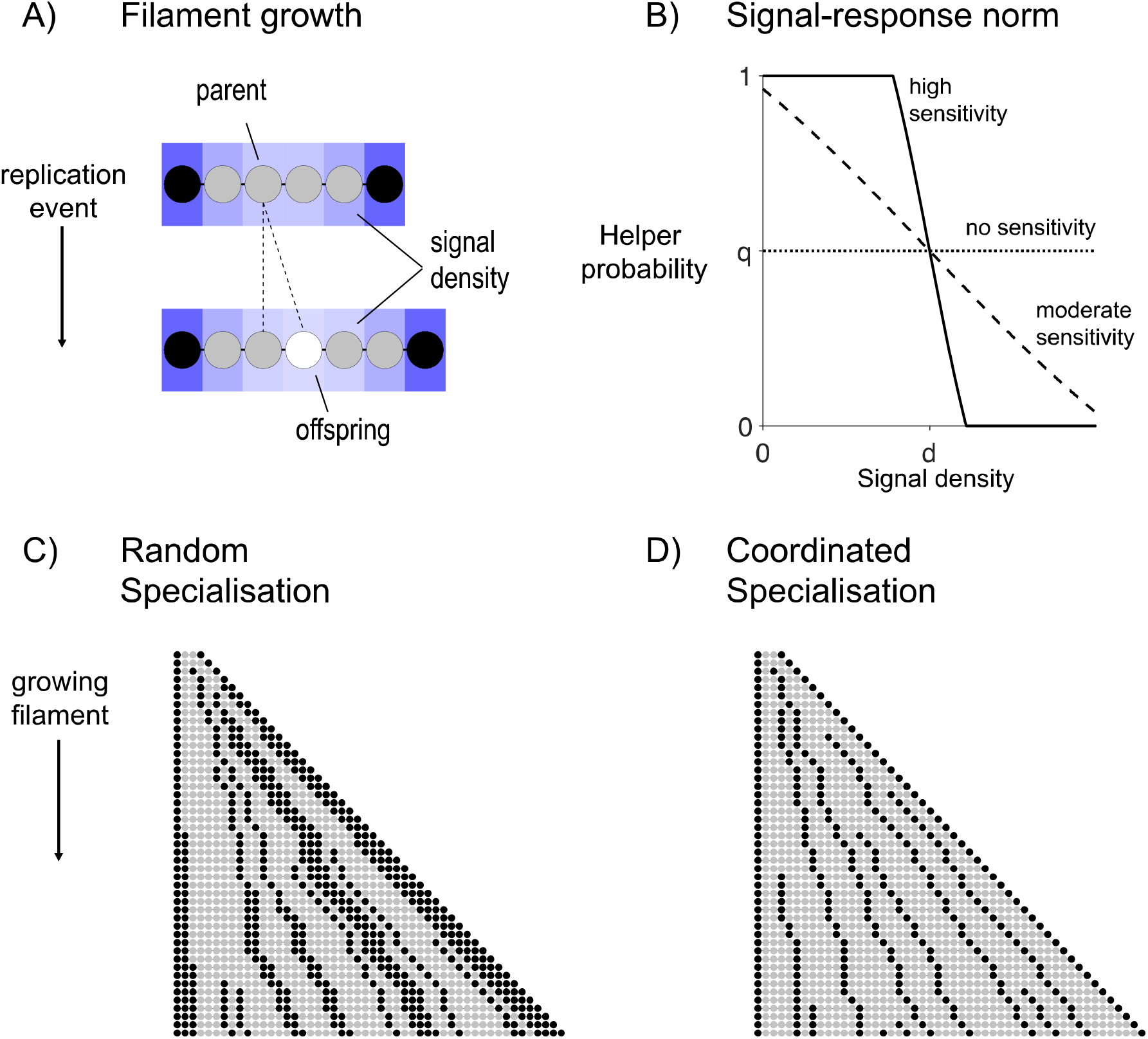
Division of labour in a cyanobacteria filament. (A) Black cells represent helpers, and grey cells reproductives. When a reproductive replicates, the parent cell produces an offspring cell (white cell) to one side of itself along the filament. The blue shading shows the density of the signal molecule produced by the helpers as it diffuses along the filament. (B) When an offspring cell is sensitive to the signal (**v** > 0), a greater (lesser) signal density will decrease (increase) the probability that it becomes a helper **(q = 0. 5, d = 5, v = 0, 0.2, 1)**. (C) A simulated example of a filament growing that employs random specialisation **(q = 0.33, s = 0, d = 0, and v = 0)**. (D) A simulated example of a filament growing that employs coordinated specialisation (**q = 0.33**, **s = 0.1**, d = 1 and **v = 1.5**) (Supplementary Section G). The helper cells (black) are more evenly spaced out (less clumped)_with coordination specialisation, compared to random specialisation.

### Division of labour in a cyanobacteria filament

We then developed a more mechanistically detailed model of a growing cyanobacteria filament to investigate the impact of within-group spatial structure (Supplementary Section G). When there is insufficient fixed nitrogen (N_2_) in the environment, some cyanobacteria species will facultatively divide labour between reproductive cells (autotrophs) that photosynthesise light and sterile helper cells (heterocysts) that fix and secrete environmental N_2_ (Fig. 1B)^9,52,53^. The fixed N_2_ diffuses along the filament where it is used by reproductives to grow and produce new cells. Division of labour in cyanobacteria is a canonical example of coordinated specialisation as helpers produce a variety of signalling molecules that diffuse along the filament to ensure that a regular pattern of phenotypes develops (Fig. 1B)^9,52,53^. Previous models of cyanobacteria focused on determining the signalling and regulatory network required to recreate the exact pattern of heterocysts along the filament^52,54–58^.

Cyanobacteria spores (hormogonium) tend to contain multiple cells^9,52^. In order to consider the case where cooperation is essential, we assume that each filament begins as a clonal sequence of two reproductives (R) and two helpers (H) in the arrangement H-R-R-H. In Supplementary Section G.4, we find that same qualitative results for the alternative assumption where all spore cells are reproductive (R-R-R-R). Over time, the number of cells in the filament increases as reproductives grow and divide by binary fission to produce within-filament offspring cells, which become either helpers or reproductives (Fig. 4A). The group life cycle ends when the filament has reached a maximum size of *L* cells. At this time, the reproductives in the filament produce dispersing spores that found filaments in the next generation of the group life cycle and all remaining cells die (non-overlapping generations) ^9^.

Reproductives grow over time by absorbing fixed N_2_, until they reach a critical size for cellular replication. Each reproductive receives fixed N_2_ from the abiotic environment at a rate of *ϕ* ≥ 0 units of fixed N_2_ per unit time (uniform background density of fixed N_2_)^58^. In addition, each helper in the filament produces fixed N_2_, at a maximum rate of units of 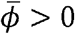 fixed N_2_ per unit time. We assume that the fixed N_2_ produced by a helper disperses across the filament with a diffusion factor, 0< *η* ≤ 1, where limited diffusivity (small *η*) means that only reproductives near the helper benefit from the fixed N_2_ it produces and high diffusivity (large *η*) means that even distant reproductives along the filament benefit. For the purposes of a focused analysis on reproductive division of labour, we ignore other forms of phenotypic heterogeneity that cyanobacteria filaments may engage in, such as the production of ATP for the group by autotrophs (non-reproductive division of labour) and the formation of persistor cells in some environments (bet-hedging)^1,53^.

Upon replication, whether a new cell becomes a helper or a reproductive depends on four evolutionary traits that jointly determine the extent of division of labour and coordination in the filament (*q, s, d*, and *v*; Fig. 4B). The baseline probability (0≤ *q* ≤ 1) is the underlying probability that a cell becomes a helper in the absence of coordination. The level of signalling (0≤ *s* ≤ 1) is the fraction of resources that a helper commits to the production and secretion of signalling molecules. The signalling molecules produced by a helper disperses along the filament with a diffusivity that we assume is distinct from the N_2_ diffusivity (Fig. 4A). The local density of signalling molecules allows new cells to estimate how close they are to a helper, or how many helpers there may be nearby.

Whether and how the new cell responds to the signal depends on the response sensitivity (*v* ≥ 0) and the response threshold (*d* ≥ 0; Fig. 4B). If *v* = 0, then a new cell is insensitive to the signal and adopts the helper phenotype with the baseline probability *q* (random specialisation). If the new cell is sensitive to the signal (*v*> 0) then a local signal density that is greater than the response threshold, *d*, will lead the cell to being less likely to adopt the helper phenotype (Fig. 4B). A higher signal density than the threshold produces the opposite effect. As sensitivity increases (higher *v*), the response to the signal becomes more deterministic (Fig. 4B).

Increasing levels of coordination (higher *v* and *s*), allows for a more precise patterning of helpers and reproductives in the filament (compare Figs. 4C and 4D). However, we assume that increased coordination is costly. First, as helpers produce more signalling molecules (higher *s*), they can produce proportionally less fixed N_2_. Second, new cells that are more sensitive to the local density of the signalling molecule (higher *v*) incur a more severe time-delay before they can specialise, such that reproductives ultimately take longer to reach the critical size of replication.

Cyanobacteria filaments employ such a signalling system and do not simply use the local density of fixed N_2_ as a cue. A possible reason for this is that signalling molecules could be fast to produce and secrete and thus coordination can occur even before helpers begin to fix N_2_^57^. Furthermore, using a dedicated signal could be more reliable than one based on fixed N_2_ density alone, which might be biased by transient fluctuations in the background level of fixed N_2_ (*ϕ*).

### Simulations

We simulated an evolving population to estimate the strategy that is favoured by natural selection in different scenarios (*q*\*, *s*\*, *d*\*, *v*\*) (Supplementary Section G). We started with a uniform population that specialises randomly (*s* = *d* = *v* = 0), and allowed the helper probability (*q*) to mutate and evolve for 500 generations, until an approximate equilibrium was reached. We then held the baseline helper probability (*q*) fixed and allowed the coordination traits (*s, d* and *v*) to mutate and evolve for 3500 generations. Each generation, the mutant strategy successfully replaces the resident strategy if it has a higher estimated average fitness. We calculate the fitness of individual filaments as the summed fecundity of reproductives in the last generation of the group life cycle, divided by the amount of time that it took the filament to grow to L cells. The separate phases of the evolutionary simulation facilitate cleaner convergence of trait-values, with an equilibrium generally being reached within 100-200 generations (Supplementary Fig. S5).

We found that the degree to which specialising cells evolve to coordinate can depend on social and environmental factors. In particular, both a lower background density of fixed N_2_ (small *ϕ*) and more limited diffusion of fixed N_2_ along the filament (smaller *η*) lead to the evolution of higher signalling levels (larger *s*\*; Fig. 5A) and higher response sensitivities (larger *v*\*; Fig. 5B). This produced filaments with more precise allocation of labour across the filament (Fig. 5E). We quantify the extent of coordination by dividing the variance in the number of helpers in a contiguous sub-block of 10 cells by the variance that would be expected for a binomial random variable of the same mean (Supplementary Section G.4.). Higher values of the reciprocal of this ratio suggest more precise division of labour.

**Figure 5:**
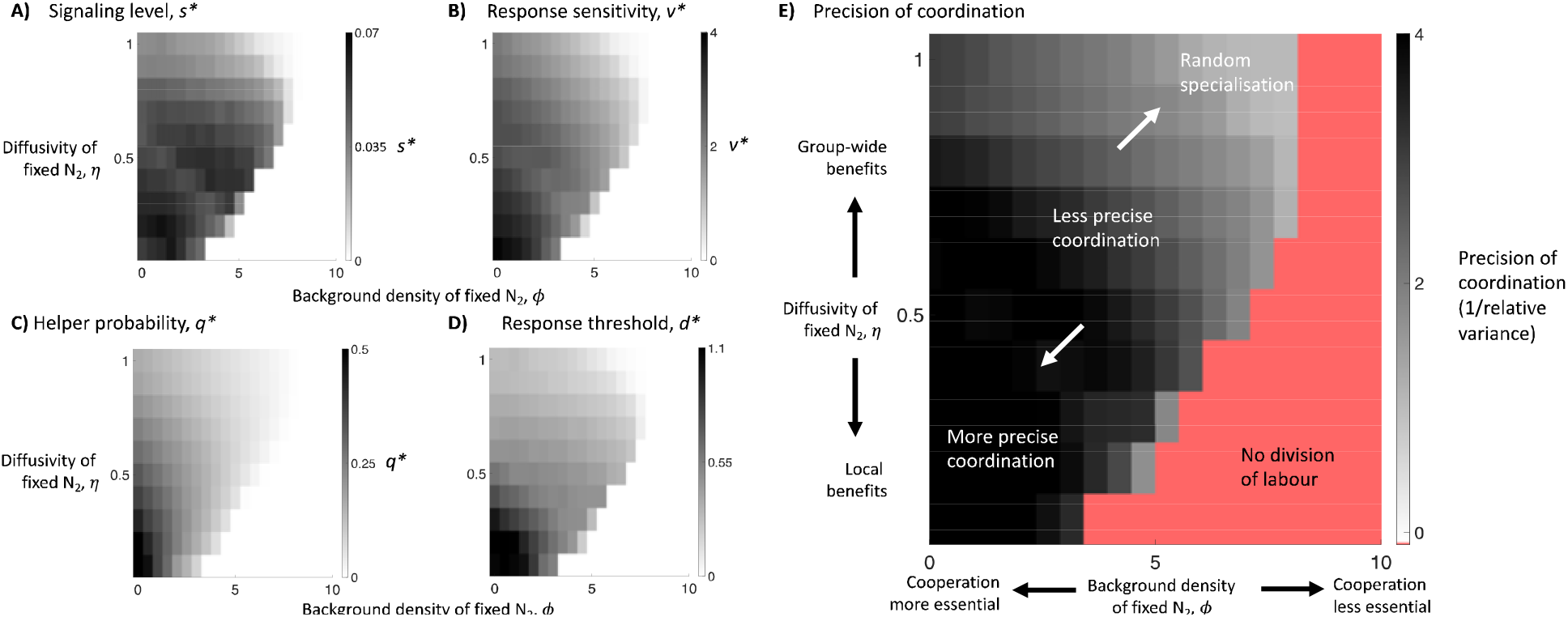
The optimal level of coordination. We present simulation results for two key factors that affect the optimal level of coordination (Supplementary Section G.4.) A lower background density of fixed ***N*_2_** (smaller ***ϕ***) and more limited diffusion of helper-fixed ***N*_2_** (smaller ***η***) favours both: (A) the evolution of a higher level of signalling (larger *s*\*); (B) a higher response sensitivity to the signal (larger *v*\*); (C) a higher baseline helper probability (larger ***q***\*); and (D) a higher response threshold (larger ***d***\*); (E) The effect of higher levels of both signalling (larger *s*\* in (A)) and response sensitivity (larger *v*\* in (B)) is that groups form with more precisely coordinated number of helpers. The precision of coordination is calculated by dividing the variance in the number of helpers in a contiguous sub-block of 10 cells relative to the variance that would be expected for a binomial random variable of the same mean (Supplementary Section G). Higher values of the reciprocal of this ratio suggest more precisely coordinated division of labour.

The predictions of our cyanobacteria model agree broadly with those of our simpler analytical model. When there is limited diffusion of helper-fixed *N*_2_ (low *η*), reproductives must depend primarily on helpers that are nearer along the filament, producing a smaller effective social group size (analogous to lower *n*). With random specialisation, a smaller social group can lead to proportions of helpers that deviate more from the optimum, increasing the benefit that can be obtained by coordination (Fig. 3). When the background density of fixed N_2_ is small (low *ϕ*), this increases the relative benefit of cooperation (analogous to higher *h*/*b*). With an increased benefit from cooperation there is a greater advantage from coordinating to produce the optimum proportion of helpers (Fig. 3). In addition, our cyanobacteria model shows how intermediate coordination can be favoured in certain scenarios (Fig. 5).

However, care is required when examining factors in mechanistic models that can have additional effects unaccounted for by their analogues in simpler models. For instance, an increase in the background density of fixed N_2_ (higher *ϕ*) means that cooperation is relatively less important (lower *h*/*b*), which we have found favours less coordination (Fig. 5). Relatively less important cooperation (lower *h*/*b*) in the mechanistic model also means that helpers may be willing to dedicate more effort to signal production (higher *s*) as there is then a relatively lower fitness cost to producing less of the public good. Another example is how helpers that produce more fixed N_2_ (larger 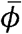) leads to cooperation that is relatively more important (higher *h*/*b*) but can also lead to larger effective social groups sizes (larger *n*) as the increased good that helpers produce can then diffuse further along the filament and benefit reproductives that are farther away.

### Spatial structure and helper clumping

Our simulations show that coordination (*s*\* > 0, *v*\* > 0) is often favoured over random specialisation ( *s*\* ≈ 0, *v*\* ≈ 0; Figs. 5A and 5B). In social groups with rigid spatial structure and local cooperation (lower *η*), an effective division of labour requires a regular distribution of helpers across the group. We hypothesized that random specialisation is particularly disadvantageous in such groups because it can lead to contiguous groups of helpers (clumps) that expand as the whole group grows (compare Figs. 4C and 4D; Supplementary Fig. S7). The helpers within these clumps can neither reproduce to break up the clump, nor are they close enough to reproductives to provide fixed N_2_. We performed additional simulations to investigate the likelihood and impact of helper clumping in growing filaments.

We found that a lower background density of fixed N_2_ (smaller *ϕ*) and more limited diffusion of fixed N_2_ (smaller *η*), leads to randomly specialising filaments with a larger average clump size (measured in number of helpers per clump; Fig. 6A), and a higher cost of clumping (measured as the slope of the best-fit line of average clump size on relative filament fitness; Fig. 6B). A higher propensity to form clumps arises because a lower background density of fixed N_2_ (smaller *ϕ*) and more limited diffusion of fixed N_2_ (smaller *η*) means new cells are more likely to become helpers (larger *q*\*; Fig. 5C). A higher cost to clumping arises in this case (smaller *ϕ* and *η*) because reproductives that are far from helpers have much lower fecundity, which increases the pressure for an even distribution of helpers. Combined, these patterns help to explain why random specialisation is disfavoured in this extreme (lower left corner of Figs. 5A, 5B, and 5E).

**Figure 6:**
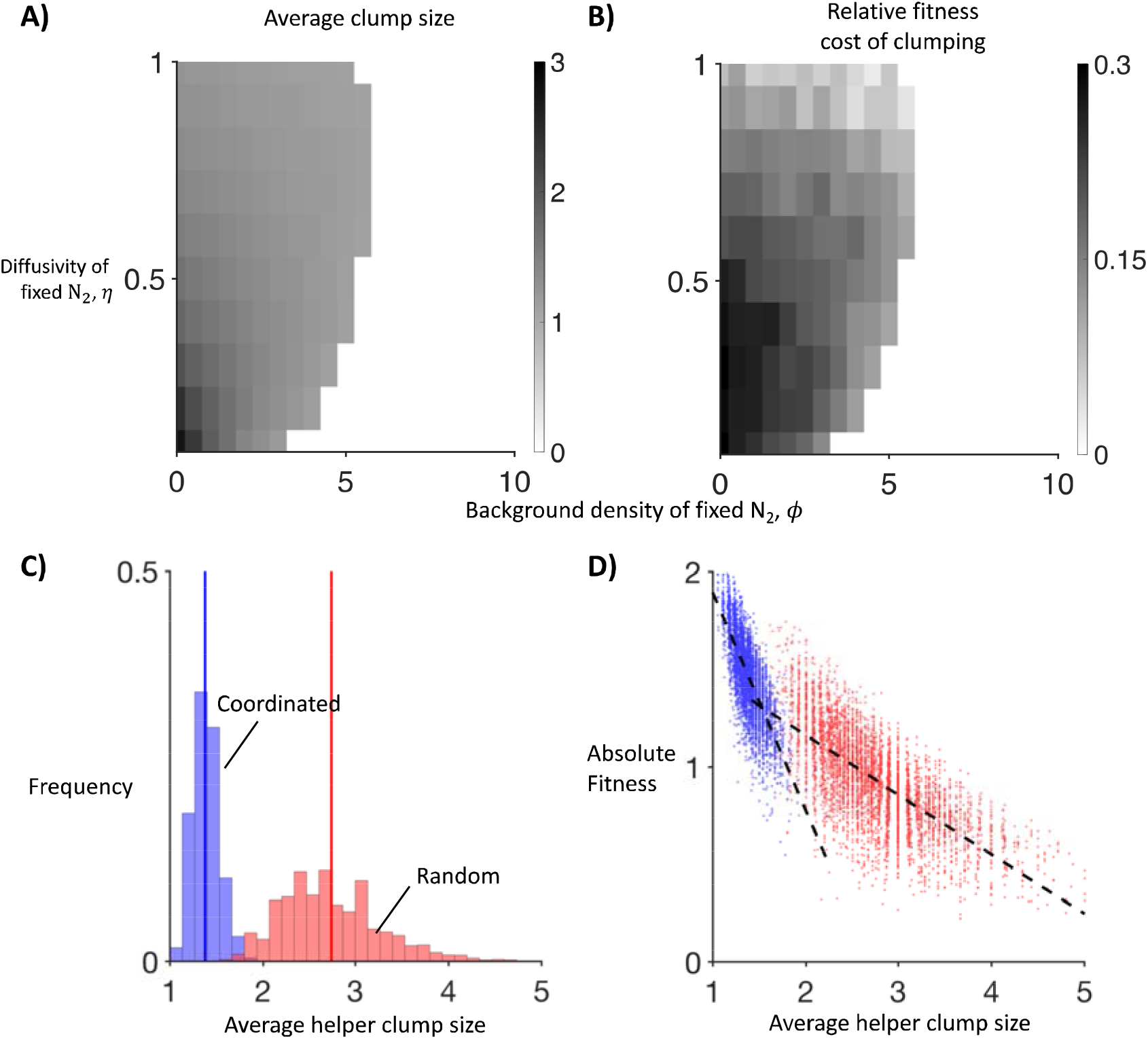
Spatial structure and helper clumping. We found that a smaller background density of fixed **N_2_** (smaller ***ϕ***) and more limited diffusion of helper-fixed **N_2_** (smaller ***η***), lead to filaments with: (A) a larger average clump size, measured as the average number of helpers per clump; and (B) a higher fitness cost of clumping, measured as the slope of the least-squares linear regression of relative fitness on average clump size. We constructed (A) & (B) by performing 1000 independent simulations of growing filaments for each parameter combination, where the trait values are set to the associated optima determined in the previous analysis (***q***\*, ***s***\*, ***d***\* and ***v***\*). We then performed 5000 independent simulations of both coordinated (blue) and random (red) filament growth at the extreme case of essential cooperation (***ϕ* = 0**) and very limited diffusion of fixed **N_2_ (η = 0.1)**. (C) Coordination leads to a dramatic reduction in average clump sizes across filaments (average clump size for random (red): 1.4 helpers and coordinated (blue): 2.7 helpers). (D) The absolute fitness cost of larger clumps is greater for coordination specialisation (blue) than for random specialisation (red) but filaments that pay the higher cost of coordination are rare. Slope of least-squares linear regression for random: −0.29 and coordinated: −1.14. Mean squared error of fit for random: 0.0181 and coordinated: 0.0297.

Focusing on the extreme case of essential cooperation (*ϕ* = 0) and very low diffusion of fixed N_2_ (*η* = 0.1), we found that coordination has two effects on clumping. Firstly, the fitness cost of clumping is more severe in coordinated filaments than for randomly specialising filaments (Fig. 6D). This occurs because coordinated helpers also invest in signalling molecules and so produce less of the public good than randomly specialised helpers, which amplifies the costs of clumping. However, secondly, coordination leads to a large reduction in the average size of clumps, and so the cost associated with larger clumps is almost never paid (Figs. 6C and 6D). Consequently, coordination ( *s*\* > 0, *v*\* > 0) can produce a substantial fitness advantage in spatial groups by decreasing the chance that costly helper clumps can form and grow.

## DISCUSSION

Our analyses provide a theoretical framework to help explain why different species of microorganisms use different mechanisms to divide labour ^2^. While testing our predictions with a formal comparative analysis would require data from more species, our predictions can help to understand the mechanisms that have evolved in well studied examples.

There are many reasons why coordinated specialisation was favoured to evolve in cyanobacteria filaments. First, cyanobacteria only divide labour when fixed N_2_ is growth-limiting and so the relative importance of cooperation is high (low *ϕ* and high *h*/*b*)^9,53,58^. Second, the fixed nitrogen produced by helpers diffuses across the filament, preferentially aiding nearby reproductives and so the effective social group size is small (low *η* and small *n*) ^9,46,59^. Third, the initial costs of coordination may have been quite small as new cells could use the local level of fixed N_2_ as a cue (low *η*) ^60^. Finally, cyanobacteria filaments have a rigid spatial structure with local benefits from cooperation and thus random specialisation could have led to the accumulation of large sterile clumps.

Colonies of *Volvox carteri* and *Dictyostelium discoideum* use coordination to divide labour, despite the fact that these groups are composed of large numbers of cells (high *n*; on the order of 1000s of cells or more) ^20,61–63^. This highlights that no single factor can fully explain empirical patterns, and that further factors not captured by simple models might be relevant in specific cases. For instance, colonies of *Volvox carteri* require a specific spatial distribution of flagella beaters across the group, which may create a strong selection pressure for coordination, analogous to the avoidance of clumps in cyanobacteria filaments. Furthermore, in some cases, details of the mechanism of division of labour are still not well understood. For instance, it is possible that there is also an initially random component to pre-stalk specialisation in *Dictyostelium*^62^.

There are multiple reasons why random specialisation would have been favoured to evolve in other well-studied species. In *Salmonella enterica*, the self-sacrificing helper cells provide a competitive advantage that eliminates other microbes but is not “essential” to the replication of *Salmonella* cells (lower *h*/*b*)^12,13^. Further, the benefits of cooperation are provided to all cells in the co-infection (*η* = 1) and so the effective social group size is reasonably large (higher *n*). Finally, *Salmonella* pathogens do not have a rigid spatial structure and so there is no scope for the accumulation of growing helper clumps as for cyanobacteria filaments. In *Bacillus subtilis*, a subset of cells become helpers that produce and secrete protein degrading proteases^43^. However, these helper cells are not sterile and so the consequence of deviating from the optimal caste ratios is reduced (Supplementary Section E.4).

To conclude, most previous work on phenotypic heterogeneity has tended to be either mechanistic, focusing on how different phenotypes are produced (caste determination), or evolutionary, focusing on why heterogeneity is favoured in the first place ^1–6,8,11,15,23–28,30,46,62,64–69^. We have used evolutionary models to explain the broader question of why different mechanisms are used in different species ^2,3,12,23–25^. Focusing on reproductive division of labour in microorganisms, we have shown that coordinated specialisation is more likely to be favoured over random specialisation in small groups, when relative coordination costs are low, and when there are larger fitness costs to deviating from optimal caste ratios. We have also shown how these patterns can hold in groups with spatial structure, where there can be a large pressure for an even distribution of phenotypes. These results identify social and environmental factors that could help to explain the distribution of mechanisms to produce phenotypic heterogeneity that have been observed in bacteria, other microbes, and beyond. Aside from microorganisms, our results also suggest a hypothesis for why random caste determination has not been widely observed in animal societies. During the initial evolution of complex animal societies, group sizes were likely to be small and the relative costs of coordination might have been minor compared to each individual’s day-to-day organismal metabolic expenditure.

## DATA AND CODE AVAILABILITY

All simulated data was generated using C and Matlab. The codes and generated data used for this study are available at: https://github.com/mingpapilio/Codes_DOL_Mechanisms.

## ACKNOWLEGEMENTS

We thank: Anna Dewar, Kevin Foster, Andy Gardner, Ashleigh Griffin, Asher Leeks, Matishalin Patel, Tom Scott, and Daniel Unterwegger for their helpful comments and suggestions; St. John’s College, Oxford (GAC), the French National Research Agency (ANR) under the Investments for the Future (Investissements d’Avenir) program (grant ANR-17-EURE-0010 to the IAST; JP) and the ERC (Horizon 2020 Advanced Grant 834164; SAW) for funding. The authors would like to acknowledge the use of the University of Oxford Advanced Research Computing (ARC) facility in carrying out this work. (http://dx.doi.org/10.5281/zenodo.22558).

## AUTHOR CONTRIBUTIONS

G.A.C., S.A.W., and J.P. conceived the study. G.A.C. and J.P. and designed and analysed the analytical models, G.A.C. and M.L. designed and analysed the simulation models. G.A.C. and S.A.W. wrote the first draft. All authors contributed toward writing the final manuscript.

## SUPPLEMENTARY INFROMATION

A. Overview

B. Labour dividers and their fitness

C. Linear public goods

D. Alternative modelling assumptions

E. Alternative forms of cooperation

F. The optimal level of coordination

G. Dividing labour in a cyanobacteria filament

Supplementary Figure 1. Alternative modelling assumptions

Supplementary Figure 2. Non-linear public goods

Supplementary Figure 3. Alternative biological scenarios

Supplementary Figure 4. The optimal level of coordination

Supplementary Figure 5: Convergence to optimal trait values

Supplementary Figure 6. Simulation results for alternate starting conditions

Supplementary Figure 7. Growing groups produce larger helper clumps

Supplementary Table 1: Cyanobacteria model

