## Supplementary Information for "The evolution of mechanisms to produce phenotypic heterogeneity in microorganisms"

### The evolution of mechanisms to produce phenotypic heterogeneity in microorganisms
### Supporting Information

Guy A. Cooper*[1,2], Ming Liu†[2], Jorge Peña‡[3], and Stuart A. West§[2]

[1]St. John's College, Oxford, United Kingdom
[2]Department of Zoology, University of Oxford, Oxford, United Kingdom
[3]Institute for Advanced Study in Toulouse, University of Toulouse Capitole, Toulouse, France

### A Overview

This manuscript is organized as follows. Section B defines the strategies of fully coordinated specialisation and random specialisation, and characterises their fitness. Section C derives the main analytical result of the main text (Equation 5), that is, the condition for fully coordinated specialisation to be favoured over random specialisation when the underlying division of labour is modelled as a linear public goods game. Section D re-examines key assumptions and approximations that simplified the analysis of the public goods model. Section E considers several models for alternative biological systems leading to different kinds of public goods. Both sections D and E demonstrate the robustness and generality of our results when relaxing our assumptions and approximations. Section F gives details on the simulation model that examines the possibility of intermediate coordination. Section G gives further details on our division of labour in cyanobacteria simulations (as presented in Fig. 4 and 5 of the main text).

### B Labour dividers and their fitness

We assume that a single individual arrives on an empty patch and, through a fixed series of replications, forms a clonal group of $n$ individuals that is composed of $k$ sterile helpers and $n-k$ pure reproductives, where $k \in \mathcal{N} = \{0, 1, 2, \ldots, n\}$. The fecundity (fitness) of the group in the absence of mechanism costs, $g_{k,n}$, is measured by the per capita number of offspring that would disperse at the end of the group life cycle. This is given by:

$$g_{k,n} = \frac{(n-k) f_{k,n}}{n}, \tag{S1}$$

where $f_{k,n}$ is the fecundity of each reproductive in the group in the absence of fecundity costs. We assume that $f_{k,n}$ depends only on the proportion of helpers in the group, $p = k/n$, with $p \in \mathcal{P} = \left\{0, \frac{1}{n}, \frac{2}{n}, \ldots, \frac{n-1}{n}, 1\right\}$. Thus, we can write $f_{k,n} = F(p)$, where $F$ is a real function. We further assume that $F$ is increasing on the interval $[0,1]$, that is, the fecundity of each reproductive is increasing in the proportion of helpers in the group. We can then rewrite (S1) as

$$g_{k,n} = (1 - k/n)\, F\,(k/n) = G(k/n), \tag{S2}$$

where we have defined

$$G(p) = (1-p)F(p). \tag{S3}$$

#### B.1 Fully coordinated specialisation

With fully coordinated specialisation (C), we assume that some mechanism, such as signalling between cells, ensures that groups always form with the optimal proportion of helpers, $p^*$, where

$$p^* = \frac{k^*}{n}, \text{ with } k^* = \arg\max_{k \in \mathcal{N}} g_{k,n}. \tag{S4}$$

We posit that the fitness of a group of coordinated specialisers is given by:

$$w_{\mathrm{C}}(p^*) = (1 - c_{\mathrm{C}}) \max_{k \in \mathcal{N}} g_{k,n} = (1 - c_{\mathrm{C}}) g_{k^*,n} = (1 - c_{\mathrm{C}}) G(p^*), \tag{S5}$$

where $0 \le c_{\mathrm{C}} \le 1$ is the metabolic cost of coordination. We make no further assumptions on the functional form of $c_{\mathrm{C}}$ although we note that it could in principle depend on other model parameters, such as group size $n$.

#### B.2 Random specialisation

With random specialisation (R), each individual in the group independently becomes a helper with a given probability $q$, and a reproductive otherwise. Hence, the number of helpers in the group, that we denote by $K$, is a binomial random variable with parameters $n$ and $q$ (i.e., $K \sim \text{Binomial}(n,q)$). In the following it will also be convenient to write $Q = K/n$ for the random variable giving the proportion of helpers in the group. Within this framework, the expected fitness of a group of random specialisers is simply given by

$$w_{\mathrm{R}}(q) = \sum_{k=0}^{n} \binom{n}{k} q^k \,(1-q)^{n-k}\, (1 - c_{\mathrm{R}}) g_{k,n}, \tag{S6}$$

where $0 \le c_{\mathrm{R}} \le 1$ is the metabolic cost of random specialisation, which we assume is independent of the number of helpers $k$.

### C Linear public goods

Our main model assumes that the fecundity function $F$ is linear in $p$ and given by

$$F(p) = b + hp, \tag{S7}$$

where $b \geq 0$ is a parameter that quantifies the baseline fecundity of reproductives in the absence of cooperation and $h > 0$ is the scale of the benefits from increased cooperation (more helpers). If there is no baseline fecundity, $b = 0$, and we say that cooperation by helpers is *essential* (i.e., the fecundity of reproductives is positive if and only if there are helpers around). If $b > 0$, cooperation is *non-essential*, with a lower value of $b$ or a higher value of $h$ leading to a larger relative importance of cooperation for the fecundity of reproductives. When cooperation is non-essential ($b > 0$), the ratio $h/b$ is a useful metric for the relative importance of cooperation.

Replacing (S7) into (S3) we obtain

$$G(p) = (1 - p)(b + hp). \tag{S8}$$

To find the fitness of coordinated specialisers, we first assume that $p$ is a continuous variable, and calculate the derivative

$$G'(p) = h - b - 2hp.$$

This derivative is decreasing in $p$ (i.e., $G(p)$ is concave), and has a single root, $\hat{p}$, given by

$$\hat{p} = \frac{h - b}{2h} = \frac{1}{2}\left(1 - \frac{b}{h}\right). \tag{S9}$$

Such a root lies in the interval $(0, 1)$ if and only if $h > b$. Otherwise the maximiser of $G(p)$ (and hence the optimal allocation of helpers) is given by $\hat{p} = 0$ (i.e., it is optimal to have no helpers). To avoid this trivial scenario without division of labour, henceforth we assume that $h > b$ holds. Further, to make progress we approximate the optimal allocation of helpers, $p^*$, by $\hat{p}$. The actual optimal value $p^*$ will be a value near $\hat{p}$ but constrained by the permissible group compositions, since $p^* \in \mathcal{P}$ (cf. section D.1, where we relax the assumption that $p^* \approx \hat{p}$). When cooperation is essential ($b = 0$), $\hat{p} = 1/2$. When cooperation is non-essential ($b > 0$), the approximate optimal proportion (S9) is an increasing function of $h/b$ with $\lim_{b \to 0} \hat{p} = 1/2$. An approximation to the fitness of fully coordinated specialisers, to be used below, can be obtained by letting $p^* \approx \hat{p}$ in equation (S5), so that

$$w_\mathrm{C}(p^*) = (1 - c_\mathrm{C})\, G(p^*) \approx (1 - c_\mathrm{C})\, G(\hat{p}). \tag{S10}$$

To find the fitness of random specialisers, we replace (S8) into (S6) and simplify to obtain

$$\begin{aligned}
w_\mathrm{R}(q) &= (1 - c_\mathrm{R}) \sum_{k=0}^{n} \binom{n}{k} q^k (1 - q)^{n-k} (1 - k/n)(b + hk/n) \\
&= (1 - c_\mathrm{R}) \left( b\mathrm{E}\left[1 - K/n\right] + h\mathrm{E}\left[(1 - K/n)K/n\right] \right) \\
&= (1 - c_\mathrm{R}) \left( b(1 - q) + hq(1 - q) - h\frac{q(1 - q)}{n} \right) && \text{(S11)} \\
&= (1 - c_\mathrm{R}) \left( G(q) - h\mathrm{Var}(Q) \right), && \text{(S12)}
\end{aligned}$$

where we have made use of the first two moments of the binomial distribution, $\mathrm{E}[K] = nq$, $\mathrm{E}[K^2] = nq(1 - q) + (nq)^2$, of expression (S8), and of the fact that $\mathrm{Var}(Q) = q(1 - q)/n$.

In order to determine the condition under which coordinated specialisation is favoured over random specialisation, we assume in a first step that random specialisers play the strategy $q = p^*$, so that their fitness is given by

$$w_\mathrm{R}(p^*) = (1 - c_\mathrm{R}) \left( G(p^*) - h\mathrm{Var}(P^*) \right), \tag{S13}$$

where $P^* = K^*/n$ and $K^* \sim \mathrm{Binomial}(n, p^*)$ (and hence $\mathrm{Var}(P^*) = p^*(1 - p^*)/n$). This assumption simplifies our calculations and leads to results that are qualitatively similar to those that arise from the more parsimonious assumption that random specialisers play the strategy that maximises their fitness (cf. section D.2, where we assume that random specialisers play optimally). We can evaluate the condition for coordinated specialisation to be favoured over random specialisation (i.e., when $w_{\mathrm{C}}(p^*) > w_{\mathrm{R}}(p^*)$ holds) by comparing expressions (S5) and (S13). This condition is given by

$$\mathrm{Var}(P^*)\frac{h}{G(p^*)} > \underbrace{\frac{c_{\mathrm{C}} - c_{\mathrm{R}}}{1 - c_{\mathrm{R}}}}_{\gamma}. \tag{S14}$$

The left-hand side of this inequality is the normalised fecundity benefit of switching from random specialisation to coordinated specialisation, and the right hand side of the inequality ($\gamma$) is the normalised relative change in metabolic costs paid from doing so. Inequality (S14) shows that the fecundity benefit of coordination over random specialisation can be decomposed into a measure of the deviation from the optimal allocation of labour, $\mathrm{Var}\,(P^*)$, and a quantity that captures the relative cost of deviating from the optimal proportion of helpers, $h/G(p^*)$.

To obtain a simple expression of condition (S14) in terms of our parameters $n$, $b$ and $h$, we approximate $p^*$ by $\hat{p}$ as given in (S9) to obtain

$$\mathrm{Var}(P^*) \approx \frac{(h-b)(h+b)}{4nh^2}, \tag{S15}$$

$$\frac{h}{G(p^*)} \approx \frac{4h^2}{(h+b)^2}. \tag{S16}$$

With these approximations, condition (S14) becomes

$$\frac{h-b}{n(h+b)} > \gamma. \tag{S17}$$

Note that the left hand side is increasing in the benefits of cooperation $h$ and decreasing in group size $n$ and the baseline fecundity $b$. Since $G(p^*)$ is (approximately) independent of $n$, we can say that the effect of increasing the group size acts primarily on the deviation of groups from the optimal proportion of helpers, $\mathrm{Var}\,(P^*)$. In contrast, a smaller baseline fecundity (lower $b$) or more benefits from cooperation (larger $h$) both (i) push $p^*$ closer to $1/2$ (which in turns increases the variance $\mathrm{Var}(P^*)$) and (ii) increases the cost of deviation (larger $h/G(p^*)$) and thus acts via both factors.

### D Alternative modelling assumptions

#### D.1 Discrete proportions of helpers

In section C, we approximated the optimal proportion of helpers in coordinated groups as a continuous variable $p^* \approx \hat{p} \in [0, 1]$. In reality this quantity is discrete as it is given by $p^* = k^*/n$ and there can only be an integer number of helpers in the group (i.e., since $k^* \in \mathcal{N}$, then $p^* \in \mathcal{P}$).

Here, we show that this assumption is relatively innocuous. First, it is clear that $p^*$ tends to $\hat{p}$ as $n$ grows large (Fig. S1A). Second, we find that even for relatively small group size $n$ the predictions of the model are relatively independent of this assumption. To show this, we repeat our analysis without making the continuous approximation. That is, we evaluate $w_{\mathrm{C}}(p^*) > w_{\mathrm{R}}(p^*)$ where $p^*$ is not approximated as $\hat{p}$ but is calculated numerically for each combination of parameters considered, so that $p^* \in \mathcal{P}$. The results of this analysis are shown in Fig. S1B. As expected, we find the broad results are similar to those of the continuous treatment, although we now find that in some cases division of labour is not favoured.

#### D.2 Optimal random specialisers

In section C, we assumed that the probability of becoming a helper, $q$, was approximately equal to the optimal proportion of helpers (i.e., we let $q = \hat{p} \approx p^*$) (Fig. S1A). A more parsimonious assumption is that the probability of being a helper in random specialisers is the one optimising fitness, so that random specialisers play strategy $q = q^*$, with

$$q^* = \arg\max_{0 \leq q \leq 1} w_{\mathrm{R}}(q). \tag{S18}$$

This alternative modelling assumption makes the fitness of random specialisers larger (as $w_{\mathrm{R}}(q^*) \geq w_{\mathrm{R}}(p^*)$ necessarily holds) and thus will make, all else being equal, random specialisation more likely to be favoured over coordinated specialisation.

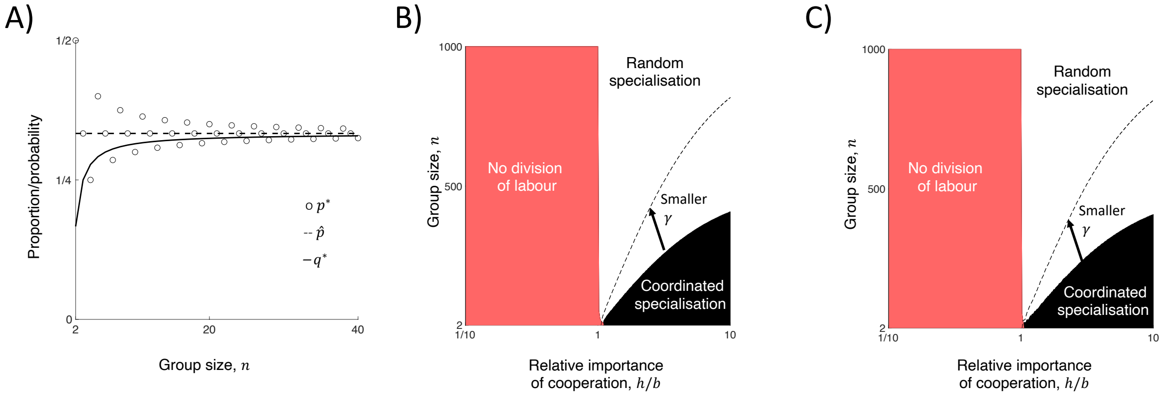

Figure S1: **Alternative modelling assumptions**. A) Optimal proportion of helpers ($p^*$; *open circles*), its continuous variable approximation ($\hat{p}$; *dashed line*), and the probability optimising fitness for random specialisers ($q^*$; *solid line*) as a function of group size $n$. In the model presented in the main text, we assumed that the probability that random specialisers adopt a helper role was equal to the optimal proportion of helpers in the group ($p^*$) as opposed to the probability that optimises fitness for random specialisers ($q^*$). In turn, we approximated this optimal proportion of helpers by a continuous variable $\hat{p}$. These three quantities converge to the same value as group size ($n$) increases. Here, $b = 1/4$ and $h = 3/4$, leading to $\hat{p} = (1/2)(1 - b/h) = 1/3$. B) Favoured mechanism (*coordinated specialisation* if $w_{\mathrm{C}}(p^*) > w_{\mathrm{R}}(p^*)$; *random specialisation* if $w_{\mathrm{C}}(p^*) < w_{\mathrm{R}}(p^*)$) without approximating $p^*$ by $\hat{p}$ (as in the main text). Instead, we evaluate the optimal value $p^*$ numerically for each parameter combination considered. The *red* region indicates where division of labour is not favoured, that is, when the optimal proportion of helpers is zero. C) Favoured mechanism (*coordinated specialisation* if $w_{\mathrm{C}}(p^*) > w_{\mathrm{R}}(p^*)$; *random specialisation* if $w_{\mathrm{C}}(p^*) < w_{\mathrm{R}}(p^*)$), while allowing for the probability that random specialisers adopt a helper role to evolve to the value that maximises the fitness of random specialisers ($q^*$). In both scenarios B) and C), we find that smaller group sizes (smaller $n$), relatively more important cooperation (higher $h/b$), and smaller relative metabolic costs of coordination (smaller $\gamma$) favour coordinated specialisation, in agreement with the results presented in the main text. In both cases, the relative importance of cooperation, $h/b$, is plotted on a log scale. In both cases we set $\gamma$ to either $1 \times 10^{-3}$ (smaller relative cost of coordination) or $2 \times 10^{-3}$ (larger relative cost of coordination).

To find $q^*$ for the linear public goods model, we take the derivative of (S11) with respect to $q$:

$$w'_{\mathrm{R}}(q) = (1 - c_R)\left(-b + h\frac{n-1}{n} - 2h\frac{n-1}{n}q\right). \tag{S19}$$

This derivative is decreasing in $q$ (hence $w_{\mathrm{R}}(q)$ is concave) and has a single root $q^*$ given by

$$q^* = \frac{h(n-1) - bn}{2h(n-1)} = \frac{1}{2}\left(1 - \frac{n}{n-1}\frac{b}{h}\right). \tag{S20}$$

Such root lies in the interval $(0, 1)$ if $w'_{\mathrm{R}}(0) = h(n-1)/n - b > 0$ holds, or equivalently, if $h/b > n/(n-1)$ holds, which we assume in the following. Expression (S20) is increasing in both $n$ and $h/b$. We also note that

$$\hat{p} - q^* = \frac{b}{2h(n-1)} \geq 0, \tag{S21}$$

and hence (i) for essential cooperation ($b = 0$), $\hat{p} = q^*$ ($b = 0$), (ii) for non-essential cooperation ($b > 0$), $\hat{p}$ always overestimates $q^*$, but that (ii) the difference between the two values is inversely proportional to $n$ and goes to zero as $n$ grows large. Thus, $\hat{p}$ approximates $q^*$ relatively well for relatively large $n$, which justifies our use of $\hat{p}$ as an approximation in section C (see Fig. S1A).

We further note that the derivative (S19) can be understood as a selection gradient on $q$. As such selection gradient is decreasing in $q$, the point $q^*$ is not only a fitness maximum, but also the value to which evolution by small-step mutations would eventually approach, that is $q^*$ is a convergence stable strategy [2, 6].

We present numerical results in Fig. S1C, evaluating the condition $w_{\mathrm{C}}(p^*) > w_{\mathrm{R}}(q^*)$ to determine when coordinated specialisation is favoured over random specialisation. Overall, we find similar qualitative results hold as in the simplified model.

##### D.3 Non-linear benefits to cooperation

In the main model, we assumed that reproductive fecundity depends linearly on the proportion of helpers in the group. Here, we consider the possibility that there is a non-linear dependence on the proportion of helpers by modelling reproductive fecundity with the sequence

$$f_{k,n} = b + h\,(k/n)^{\nu}, \tag{S22}$$

where $\nu > 0$ is a parameter controlling the shape of the return from an increasing proportion of helpers. When $\nu < 1$, there is a large initial return from the addition of the first helper, followed by a decelerating rate of return as the proportion of helpers increases (i.e., $f_{k,n}$ is concave; see Fig. S2A). When $\nu > 1$ there is a small initial return from the addition of the first helper followed by an accelerating rate of return as the proportion of helpers increases (i.e., $f_{k,n}$ is convex; see Fig. S2B). In all cases, (S22) is monotonically increasing from zero to one.

We considered the parameter discretisation, $n \in \{2, 4, \ldots, 38, 40\}$ and $h/b \in \{1, 1 + 1 \times 9/24, \ldots, 1 + 23 \times 9/24, 1\}$ and for each combination of parameters, we solved numerically for the optimal proportion of helpers, $p^*$, that maximises $G(p)$. For each combination of parameters, we then determined whether coordinated specialisation is favoured over random specialisation ($w_{\mathrm{C}}(p^*) > w_{\mathrm{R}}(p^*)$), to ascertain whether accelerating ($\nu > 1$), or decelerating ($\nu < 1$) returns affect the predictions of our analysis. We find that the same broad results hold as in the linear public goods game (see Fig. S2C and S2D). We further find that accelerating benefits, in which there is a low initial return from helpers, can only lead to division of labour being favoured for a sufficiently high relative importance of cooperation (Fig. S2D).

#### E Alternative forms of cooperation

In the main model, we assumed that the fecundity of reproductives depends on the proportion of sterile helpers in the group, $k/n$. Here, we consider the conceptual underpinnings of this assumption (section E.1) and analyse alternative scenarios where reproductive fecundity depends on the number of helpers in the group, $k$ (section E.2), or on the ratio of helpers to reproductives, $k/(n-k)$ (section E.3). We also analyse a model in which helpers may also reproduce, albeit with diminished fecundity (section E.4). Further, our main model assumed that division of labour occurs only when groups reach their maximal size. Here we consider an extention of the model in which division of labour occurs in every generation of group growth (section E.5).

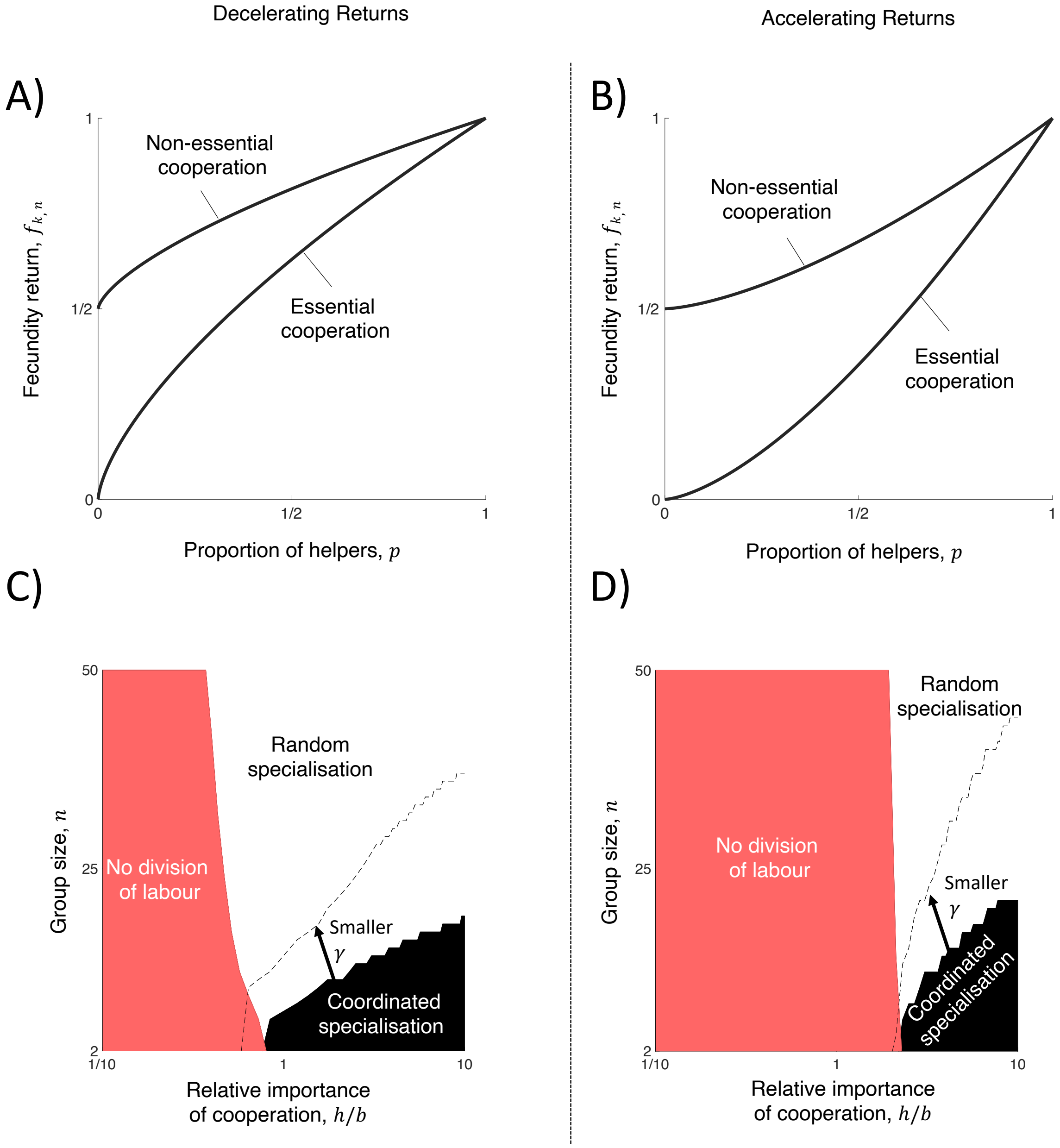

Figure S2: **Non-linear public goods**. We consider an extension of the model allowing the fecundity of reproductives to depend non-linearly on the proportion of helpers in the group. (A) If $\nu < 1$, then the presence of helpers provides an initially large increase in the fecundity of reproductives but then only provides benefits at a decreasing rate as the proportion of helpers goes up (decelerating cooperation; here $\nu = 2/3$). (B) If $\nu > 1$, then the presence of helpers provides no initial increase in the fecundity of reproductives but then provides benefits at an increasing rate as the proportion of helpers goes up (accelerating cooperation; here $\nu = 3/2$). In both A) and B) we have set $b = h = 1/2$ for non-essential cooperation and $b = 0$ and $h = 1$ for essential cooperation. (C and D) Favoured mechanism (*coordinated specialisation* if $w_{\mathrm{C}}(p^*) > w_{\mathrm{R}}(p^*)$; *random specialisation* if $w_{\mathrm{C}}(p^*) < w_{\mathrm{R}}(p^*)$) with fecundity sequence given by Eq. (S22) and either $\nu = 3/4$ (decelerating cooperation; panel C) or $\nu = 4/3$ (accelerating cooperation; panel D). In both scenarios, we find that small group sizes (lower $n$), relatively more important cooperation (higher $h/b$) and a smaller relative metabolic cost of coordination (lower $\gamma$) favour coordinated specialisation. In both cases, the relative importance of cooperation, $h/b$, is plotted on a log scale. We set $\gamma$ either to $= 0.02$ (lower relative cost of coordination) or $0.04$ (higher relative cost of coordination). The *red* region shows where division of labour is not favoured, that is, when the optimal proportion of helpers is zero.

#### E.1 Reproductives benefit from the proportion of helpers

There are two key scenarios where reproductive fecundity can be modelled as depending on the proportion of helpers in the group, $k/n$.

First, the benefits provided by helpers may constitute a good that is consumed by its beneficiaries (sometimes termed a congestible or rivalrous good), wherein the benefit experienced by one individual proportionally decreases the amount of good that may benefit its neighbours [3, 5]. For instance, the secretion of an extracellular product that must be absorbed and digested in order to provide benefits (as happens in populations of *B. subtilis* and *A. cylindrica*) can be conceptualised as such a good [7, 12, 13]. In particular, if the good is non-excludible (i.e., all individuals use the benefits) then the benefits conferred to each reproductive is the total amount of help (in the linear case, approximated as the number of helpers, $k$) divided by the number of individuals in the group ($n$; reproductives and helpers alike) [5]. The fact that helpers also partake in the consumption of the public good may be considered a type of soaking [10].

Second, the amount of good that each helper provides may depend on the number of individuals in the group such that, all else being equal, larger groups entail less of a benefit to the group. For instance, helpers in *V. carteri* beat their flagella to keep the colony afloat at the optimal height in the water column for photosynthesis [9, 11]. In this case, the contribution of each helper (the degree to which it helps keep the colony at the right height) depends inversely on the size of the group as larger colonies are more difficult to keep afloat.

#### E.2 Reproductives benefit from the number of helpers

An alternative assumption is that the fecundity of reproductives depends on the number of helpers in the group, $k$. This would imply that the contribution of each helper to the total good does not depend on the size of the group and that the benefit conferred to one individual does not decrease the benefits available for another. The benefits provided by helpers in *S. enterica* infections is an example of such a good (a non-congestible or non-rivalrous good) [3, 5]. In this case, helpers trigger a host immune response that wipes out competing microbial strains so that reproductives may then proliferate without competition [1, 4]. Thus, the competitive advantage afforded by the host immune response is not used up or depleted by any of its beneficiaries, unless there are so few niches vacated that reproductives then compete with one another.

We would like to know whether we recover similar results as the ones we obtained assuming that reproductives benefit from the proportion of helpers in an alternative model in which the fecundity of reproductives depends on the number of helpers in the group, $k$. To this end we posit the following specific sequence for reproductive fecundity:

$$f_{k,n} = b + hk.$$

Replacing this expression into (S3) and by some abuse of notation (as now $G$ depends explicitly on the group size $n$), we obtain

$$G(p) = (1-p)(b + hnp). \tag{S23}$$

Treating $p$ as a continuous variable, we calculate the first derivative of $G$ with respect to $p$ as

$$G'(p) = hn - b - 2hnp.$$

This derivative is decreasing in $p$ (i.e., $G(p)$ is concave), and has a single root $\hat{p}$ given by

$$\hat{p} = \frac{hn-b}{2hn} = \frac{1}{2}\left(1 - \frac{1}{n}\frac{b}{h}\right). \tag{S24}$$

Such root lies in the interval $(0,1)$ if $h/b > 1/n$, which we assume henceforth.

Using arguments similar to the ones we used in section C, it follows that we can approximate the fitness of a coordinated group as

$$w_{\mathrm{C}}(p^*) = (1-c_{\mathrm{C}})G(p^*) \approx (1-c_{\mathrm{C}})G(\hat{p}),$$

while the fitness of a randomly specialising group randomising with probability $q$ can be written as

$$\begin{aligned} w_{\mathrm{R}}(q) &= (1-c_{\mathrm{R}})\sum_{k=0}^{n}\binom{n}{k}q^k(1-q)^{n-k}(1-k/n)(b+hk) \\ &= (1-c_{\mathrm{R}})\left(b + hnq - bq - hnq^2 - hq(1-q)\right) \\ &= (1-c_{\mathrm{R}})\left(G(q) - hq(1-q)\right) \\ &= (1-c_{\mathrm{R}})\left(G(q) - hn\mathrm{Var}(Q)\right). \end{aligned}$$

Assuming that random specialisers play $q = p^*$, the condition for coordinated specialisation to be favoured over random specialisation is:

$$\mathrm{Var}(P^*)\frac{h}{G(p^*)}n > \underbrace{\frac{c_\mathrm{C} - c_\mathrm{R}}{1 - c_\mathrm{R}}}_{\gamma}. \tag{S25}$$

Further endorsing our approximation $p^* \approx \hat{p}$, we have

$$\mathrm{Var}(P^*) \approx \frac{(hn - b)(hn + b)}{4n^3h^2}, \tag{S26}$$

$$\frac{h}{G(p^*)} \approx \frac{4h^2n}{(hn + b)^2}, \tag{S27}$$

and condition (S25) becomes

$$\frac{hn - b}{n(hn + b)} > \gamma. \tag{S28}$$

We present some numerical results in Fig. S3A, evaluating the condition $w_\mathrm{C}(p^*) > w_\mathrm{R}(p^*)$. As in the main model where reproductives benefit from the proportion of helpers, we find that smaller group sizes (small $n$) and relatively more important cooperation (higher $h/b$) favour coordinated specialisation. We also find that less essential forms of cooperation ($h/b < 1$) may still favour division of labour.

#### E.3 Reproductives benefit from the ratio of helpers to reproductives

In a second alternative scenario, only reproductive individuals benefit from the collective good. In this case, the fecundity of reproductives is determined by the number of helpers (approximate amount of collective good) divided by the number of reproductives (the number of beneficiaries of the good). This gives the following expression for the fecundity of a reproductive in a group of size $n$ with $k$ helpers:

$$f_{k,n} = \left(b + h\frac{k}{n-k}\right)[k < n], \tag{S29}$$

where $[k < n]$ is an Iverson bracket indicating that $f_{k,n} = 0$ if $k = n$.

Replacing this expression into (S1) and simplifying, we obtain the following formula for the fitness of a group of size $n$ with $k$ helpers:

$$g_{k,n} = \left(b + \frac{(h-b)k}{n}\right)[k < n]. \tag{S30}$$

This sequence is unimodal (first increasing, then decreasing) in $k$ if $h > b$ holds, which we assume henceforth. In this case, $k^* = n - 1$ maximises $g_{k,n}$ for all $n$, and the optimal proportion of helpers is hence given by $p^* = (n-1)/n$. It follows that the fitness for coordinated specialisers is given by

$$w_\mathrm{C}(p^*) = (1 - c_\mathrm{C})g_{k^*,n} = (1 - c_\mathrm{C})g_{n-1,n} = (1 - c_\mathrm{C})\frac{b + (n-1)h}{n}. \tag{S31}$$

In contrast, the expected fitness for random specialisers is given by:

$$\begin{aligned} w_\mathrm{R}(q) &= (1 - c_\mathrm{R})\sum_{k=0}^{n-1}\binom{n}{k}q^k(1-q)^{n-k}[b + (h-b)k/n] \\ &= (1 - c_\mathrm{R})\left\{\sum_{k=0}^{n}\binom{n}{k}q^k(1-q)^{n-k}[b + (h-b)k/n] - \binom{n}{n}q^n(1-q)^0[b + (h-b)]\right\} \\ &= (1 - c_\mathrm{R})\left(b + (h-b)q - hq^n\right). \end{aligned} \tag{S32}$$

At this point we deviate from our previous approximations and compare the fitness of coordinated specialisers and the fitness of random specialisers by evaluating the condition $w_\mathrm{C}(p^*) > w_\mathrm{R}(q^*)$, where $q^*$ is the optimal probability of becoming a helper for random specialisers. We do this because this model leads to highly asymmetric group fitness $g_{k,n}$, which select for random specialisers to strongly under weigh the probability of becoming a helper ($q^* \ll p^*$) and therefore the approximation $q^* \approx p^*$ is less accurate.

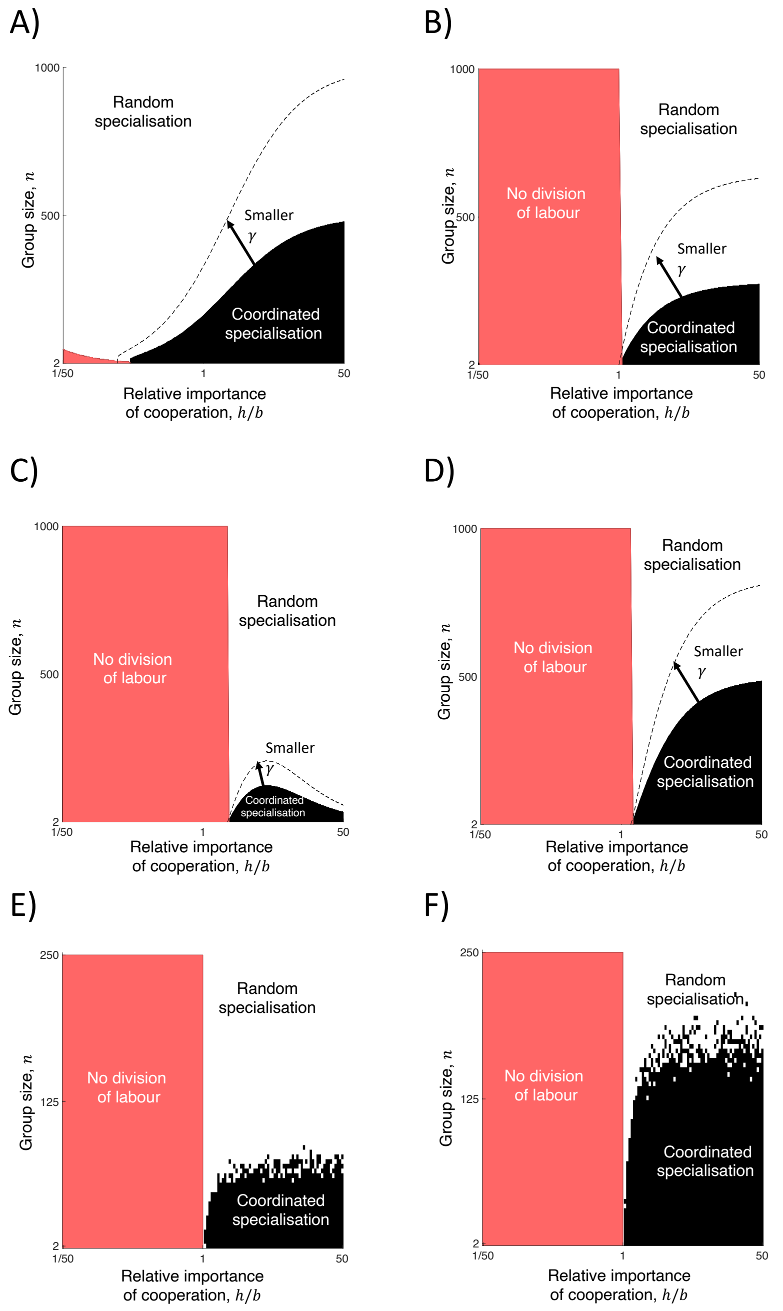

Figure S3: **Alternative biological scenarios**. We consider the relative fitness advantage of coordination over random specialisation as a function of the final size of the group $n$ and the relative importance of cooperation, $h/b$, in different biological scenarios. (A) Fecundity of reproductives depends on the number of helpers in the group. $\gamma = 1 \times 10^{-3}$ (lower relative costs of coordination) or $\gamma = 2 \times 10^{-3}$ (higher relative cost of coordination). (B) Fecundity of reproductives depends on the ratio of helpers to reproductives. $\gamma = 1 \times 10^{-2}$ (lower relative costs of coordination) or $\gamma = 2 \times 10^{-2}$ (higher relative cost of coordination). (C and D) Helpers are non-sterile but less specialised helpers spending a proportion $x$ of their resources on producing the public good (panel C: $x = 1/2$; panel D: $x = 3/4$). In both cases smaller group sizes (lower $n$), relatively more important cooperation (higher $h/b$) and a smaller relative cost of coordination ($\gamma$) favour coordinated specialisation. More specialised helpers (higher $x$) favours more coordinated specialisation. In both cases, $\alpha = 2$. For both panels C and D: $\gamma = 6 \times 10^{-4}$ (lower relative costs of coordination) or $\gamma = 1 \times 10^{-3}$ (higher relative cost of coordination). (E and F) Division of labour occurs in every generation as the group grows (panel E: $\gamma = 0.03$; panel F: $\gamma = 0.015$). Smaller group sizes (smaller $n$), relatively more important cooperation (higher $h/b$) and a smaller relative cost of coordination ($\gamma$) favour coordinated specialisation. In all cases, the relative importance of cooperation, $h/b$, is plotted on a log scale. In the *red* region division of labour is not favoured, that is, the optimal proportion of helpers is zero.

The derivative of $w_{\mathrm{R}}(q)$ (S32) with respect to $q$ is given by $w'_{\mathrm{R}}(q) = (1 - c_{\mathrm{R}})(h - b - hnq^{n-1})$. For $h > b$, this expression has a single root in the interval $(0, 1)$, given by

$$q^* = \left(\frac{h-b}{hn}\right)^{\frac{1}{n-1}}, \tag{S33}$$

which maximises $w_{\mathrm{R}}(q)$. Evaluating (S32) at $q = q^*$ and simplifying, we obtain the coordinated specialisation is favoured over random specialisation ($w_{\mathrm{C}}(p^*) > w_{\mathrm{R}}(q^*)$), when:

$$\frac{b/n + h(n-1)/n}{b + (h-b)\left(\frac{h-b}{hn}\right)^{\frac{1}{n-1}} - h\left(\frac{h-b}{hn}\right)^{\frac{n}{n-1}}} > \frac{1 - c_{\mathrm{R}}}{1 - c_{\mathrm{C}}}. \tag{S34}$$

In Fig. S3B, we graphically show when coordinated specialisation is favoured over random specialisation. We find that that smaller group sizes (small $n$) and relatively more important cooperation (higher $h/b$) favour coordinated specialisation.

In contrast to what whe found when benefits depend on the number of helpers (cf. Fig. S3A), when the benefits depend on the ratio of helpers to reproductives (Fig. S3B) and the relative importance of cooperation is high (high $h/b$), larger group sizes (large $n$) may still favour fully coordinated specialisation.

#### E.4 Reproductive division of labour with non-sterile helpers

We consider here the possibility that helpers also reproduce. This happens for instance for the subset of *Bacilus subtilis* cells that produce protein degrading proteases but still produce offspring. Let $x \in [0, 1]$ be the degree to which helpers are specialised in the production of a public good. We assume that no specialisation ($x = 0$) entails no production of public goods and no personal fecundity cost, but that increasing specialisation (higher $x$) leads to higher production of public goods at a linearly increasing personal cost to fecundity such that full specialisation ($x = 1$) means that helpers are sterile. At a given level of specialisation, the amount of public good produced by a helper is modelled as $x^\alpha$, where $\alpha > 1$ is the scale of the efficiency benefit from specialisation. Combined, these assumptions give the following fitness equation:

$$G(p) = p(1 - x)(b + hx^\alpha p) + (1 - p)(b + hx^\alpha p), \tag{S35}$$

where the first term is the per capita fitness of the group due to the fecundity of helpers and the second term is the per capita fitness of the group due to the fecundity of reproductives. To find the fitness of coordinated specialisers, we again assume that $p$ is a continuous variable, and calculate the derivative

$$G'(p) = -2hx^{\alpha+1}p - xb + hx^\alpha.$$

This derivative is decreasing in $p$ (i.e., $G(p)$ is concave), and has a single root, $\hat{p}$, given by

$$\hat{p} = \frac{hx^{\alpha-1} - b}{2hx^\alpha} = \frac{1}{2}\left(x^{-1} - x^{-\alpha}\frac{b}{h}\right). \tag{S36}$$

Such a root lies in the interval $(0, 1)$ if and only if $hx^{\alpha-1} > b$. Otherwise the maximiser of $G(p)$ (and hence the optimal allocation of helpers) is given by $\hat{p} = 0$ (i.e., it is optimal to have no helpers). Again, to make progress we approximate the optimal allocation of helpers, $p^*$, by $\hat{p}$.

Note that the approximate optimal proportion (S36): (i) is independent of $n$; (ii) is an increasing function of $h/b$ such that $\hat{p} = 1/(2x)$ when cooperation is essential ($b = 0$); and (iii) is an increasing function of the degree of helper specialisation $x$, such that $\hat{p} = (h - b)/(2h)$ when helpers are sterile ($x = 1$; this recovers Equation (S24) as a particular case).

We then obtain an approximation to the fitness of fully coordinated specialisers by letting $p^* \approx \hat{p}$, which gives

$$w_{\mathrm{C}}(p^*) = (1 - c_{\mathrm{C}})G(p^*) \approx (1 - c_{\mathrm{C}})G(\hat{p}). \tag{S37}$$

The fitness of random specialisers is given by

$$
\begin{aligned}
w_{\mathrm{R}}(q) &= (1-c_{\mathrm{R}}) \sum_{k=0}^{n} \binom{n}{k} q^k \,(1-q)^{n-k} \left[(k/n)(1-x)(b + hx^{\alpha}k/n) + (1-k/n)(b + hx^{\alpha}k/n)\right] \\
&= (1-c_{\mathrm{R}}) \left\{ b(1-x)\mathrm{E}\left[K/n\right] + hx^{\alpha}(1-x)\mathrm{E}[K^2/n^2] + b\mathrm{E}\left[(1-K/n)\right] + hx^{\alpha}\mathrm{E}[K/n - K^2/n^2] \right\} \\
&= (1-c_{\mathrm{R}}) \left\{ b(1-x)q + hx^{\alpha}(1-x)\frac{nq(1-q)+n^2q^2}{n^2} + b(1-q) + hx^{\alpha}\left[q - \frac{nq(1-q)+n^2q^2}{n^2}\right] \right\} \\
&= (1-c_{\mathrm{R}}) \left\{ b(1-x)q + hx^{\alpha}(1-x)\left[\frac{q(1-q)}{n} + q^2\right] + b(1-q) + hx^{\alpha}\left[q(1-q) - \frac{q(1-q)}{n}\right] \right\} \\
&= (1-c_{\mathrm{R}}) \left\{ q(1-x)\,(b + hx^{\alpha}q) + hx^{\alpha}(1-x)\frac{q(1-q)}{n} + (1-q)\,(b + hx^{\alpha}q) - hx^{\alpha}\frac{q(1-q)}{n} \right\} \\
&= (1-c_{\mathrm{R}}) \left\{ G(q) - hx^{\alpha+1}\frac{q(1-q)}{n} \right\} \qquad \text{(S38)} \\
&= (1-c_{\mathrm{R}}) \left( G(q) - hx^{\alpha+1}\mathrm{Var}(Q) \right), \qquad \text{(S39)}
\end{aligned}
$$

where we have made use of the first two moments of the binomial distribution, $\mathrm{E}[K] = nq$, $\mathrm{E}[K^2] = nq(1-q) + (nq)^2$, of expression (S35), and of the fact that $\mathrm{Var}(Q) = q(1-q)/n$.

We again assume that random specialisers play the strategy $q = p^*$, so that their fitness is given by

$$w_{\mathrm{R}}(p^*) = (1-c_{\mathrm{R}}) \left( G(p^*) - hx^{\alpha+1}\mathrm{Var}(P^*) \right), \qquad \text{(S40)}$$

where $P^* = K^*/n$ and $K^* \sim \mathrm{Binomial}(n, p^*)$ (and hence $\mathrm{Var}(P^*) = p^*(1-p^*)/n$). Comparing expressions (S37) and (S39), it follows that coordinated specialisation is favoured over random specialisation (i.e., $w_{\mathrm{C}}(p^*) > w_{\mathrm{R}}(p^*)$ holds) when

$$\mathrm{Var}(P^*)\frac{hx^{\alpha+1}}{G(p^*)} > \gamma. \qquad \text{(S41)}$$

Again, condition (S41) makes it explicit that the fecundity benefit of coordination over random specialisation can be decomposed into a measure of the deviation from the optimal allocation of labour, $\mathrm{Var}\,(P^*)$, and a quantity that captures the relative cost of deviating from the optimal proportion of helpers, $hx^{\alpha+1}/G(p^*)$.

To obtain a simple expression of condition (S41) in terms of our parameters $n$, $b$ and $h$, we approximate $p^*$ by $\hat{p}$ as given in (S36). Substituting this approximation, condition (S41) becomes:

$$\frac{(hx^{\alpha-1} - b)(hx^{\alpha-1}(2x-1) + b)}{n(hx^{\alpha-1} + b)^2} > \gamma. \qquad \text{(S42)}$$

It is easy to see that the left-hand side of equation S42 is decreasing in the size of the group $n$. Numerical examination reveals that the left-hand side is increasing in the degree of helper specialisation, $x$, and that it is unimodal in the relative benefits of cooperation, $h/b$, and in the efficiency benefits of specialisation, $\alpha$ (increasing and then decreasing). We show these results graphically in Fig. S3C and S3D for two different values of helper specialisation.

When helpers are less specialised (smaller $x$), more essential cooperation ($h/b$) can lead to coordinated specialisation being less likely to evolve (Fig. S3C). This occurs because more essential cooperation leads to a higher proportion of helpers ($\hat{p}$), which can increase to values greater than $1/2$ when helpers are not sterile ($x < 1$) (Equation S36). When a high proportion of non-sterile helpers is favoured ($\hat{p} \approx 1$), the expected variance in the proportion of helpers can become very small ($\mathrm{Var}(P^*) \approx 0$) and thus random groups are much less likely to deviate from the optimal proportion of helpers.

#### E.5 Reproductive division of labour as groups grow

We outline here a simulation model in which cells divide labour in every generation of the group-growth cycle. We simulate group-growth cycles using either random specialisation or coordinated specialisation in order to estimate the average fitness of each mechanism to divide labour in different scenarios. We assume that the social interactions are given by a linear public goods game as the one presented in section C. Although we investigate division of labours in groups of finite size, and for simplicity, we implicitly assume that the size of groups is large. In particular, we make the approximation $p^* \approx \hat{p}$, with $\hat{p}$ given by (S9).

For each simulation, we assume that groups start with one reproductive and that one cell is added to the group each generation of the group-growth cycle until the group has reached a total of $n$ cells. Let $t \in T = \{1, 2, \ldots, n\}$ index the generation of the group growth such that in generation $t$, there are $t$ cells in the group. Let $p_t$ be the proportion of helpers in the group at generation $t$. Since groups start with one reproductive, $p_1 = 1$.

For groups that specialise randomly with helper probability, $q$, we assume that each generation of the group growth cycle the new cell adopts a helper phenotype with probability $q$ and otherwise is a reproductive. More formally, at generation $t > 1$, we have $p_t = (p_{t-1}(t-1)+1)/t$ with probability $q$ and $p_t = (p_{t-1}(t-1))/t$ with probability $1-q$. The proportion of helpers in the final group of the life cycle, $p_n$, can take a value in the set $\{0, 1/n, \ldots, (n-1)/n\}$.

For groups that specialise with coordination, we assume that new cells adopt the phenotype that minimises the absolute difference between the proportion of helpers in the group and the optimal proportion of helpers, $p^* \approx \hat{p} = (h-b)/(2h)$ (cf. equation (S9)). More formally, in generation $t > 1$, if $|(p_{t-1}(t-1)+1)/t - p^*| > |p_{t-1}(t-1))/t - p^*|$ then the new cell adopts a reproductive phenotype, and hence $p_t = (p_{t-1}(t-1))/t$. On the other hand if $|(p_{t-1}(t-1)+1)/t - p^*| < |p_{t-1}(t-1)/t - p^*|$, then the new cell adopts a helper phenotype, and hence $p_t = (p_{t-1}(t-1)+1)/t$.

For each combination of parameter values considered, we simulated a large number of group-growth cycles (500 replications) for each mechanism in order to estimate the average fitness of each mechanism ($w_{\mathrm{C}}$ and $w_{\mathrm{R}}$). In each simulation, the fitness of the group at the end of the life cycle is calculated as the product of the group fecundities at each generation,

$$\prod_{t \in T} (1-c)(1-p_t)(b+hp_t), \tag{S43}$$

where $(1-p_t)(b+hp_t)$ is the fecundity of the group in generation $t$ (cf. equation (S8)) and we set $c = c_{\mathrm{C}}$ for coordinated groups and $c = c_{\mathrm{R}}$ for randomly specialising groups. Once again, we assume that the probability that random specialisers adopt a helper role is equal to the optimal proportion of helpers, $q = p^* \approx \hat{p}$. We also assumed that the cost of the mechanisms $c_{\mathrm{C}}$ and $c_{\mathrm{R}}$ is independent of group size, $n$, for both mechanisms.

In Fig. S3E and S3F, we show that smaller final group sizes (lower $n$), more essential cooperation (higher $h/b$) and lower relative costs of coordination (smaller $\gamma$) favour coordinated specialisation.

### F The optimal level of coordination

In the first analysis of the main text, we assumed that all individuals in a coordinated group interact with one another and so have complete information about the phenotypes of their social partners when specialising. Consequently, coordinated groups are fully coordinated and always contain the optimal proportion of helpers to reproductives, $p^*$. Here we relax this assumption and use individual based simulations to determine the optimal level of coordination.

We provide a heuristic model where increasing the number of cell-to-cell interactions within the group can lead incrementally to a more precise allocation of labour. We consider a costly trait $s \in [0,1]$ (the level of coordination) that is equal to the independent probability that any two cells in the group interact with each other. When a focal individual specialises, it adopts a helper role depending on how the proportion of helpers amongst cells that it interacts with compares to a critical threshold, $t \in [0,1]$. Variables $s$ and $t$ are co-evolving traits in our simulations; they influence both the degree to which groups are coordinated and the degree to which labour is divided (i.e., the realised proportion of helpers, $p$). As the expected connectivity of the group increases (i.e., higher $s$, more one-to-one interactions), there is a higher cost of coordination paid by the group. However, this higher cost of coordination may be offset by the increased amount of information afforded to each individual when specialising and thus an increased chance that the group ends up with the optimal proportion of helpers, $p^*$. This trade-off means that in some cases the optimal strength of coordination may in fact be intermediate ($0 < s < 1$), rather than the full coordination considered in the main analysis ($s = 1$).

In the following, we outline our individual-based model. First, we specify how the simulations are initialised. Second, given a group with a particular level of coordination ($s$), we specify how we determine which individuals interact with one another. Third, given a group with an interaction network and target proportion of helpers ($t$), we describe how the allocation of labour within the group is determined. Fourth, given the allocation of labour, we specify how the fitness of each group is determined. Finally, we describe how the individuals in the population compete globally from one generation to the next, and how mutation may affect the trait values of $s$ and $t$. Results for the evolved level of coordination and target proportion of helpers are depicted in Fig. S4.

#### F.1 Simulation initialisation

We create a population of approximately $N_{\mathrm{T}}$ cells, sorted into groups of $n$ cells. Specifically, we set the number of groups in the population as $N_{\mathrm{g}} = \lceil N_{\mathrm{T}}/n \rceil$, where $\lceil a \rceil$ is the least integer larger than or equal to $a$. Thus, the population size, $N_{\mathrm{g}}n$, is relatively constant while the number of groups increases as the group size decreases. This reduces the effect of demographic stochasticity across simulations. All individuals in the population are characterised by their trait values $s$ and $t$. We assume that all groups are founded by a single asexual individual, and thus that groups are clonal and that all individuals in the same group have the same trait values. At the beginning of the simulation, we assume that all individuals in the population have no propensity for either division of labour or coordination (i.e., $s = 0, t = 0$).

#### F.2 Coordination network

Given a group with a particular level of coordination, $s$, we determine the between-cell coordination network by constructing a random graph $G(n, s)$ where the $n$ nodes correspond to the $n$ cells in the group and where each possible edge (representing the interaction between a given pair of cells) occurs independently with probability $s$ [8]. Thus, for a given $s$, the expected number of interacting pairs is $sn(n-1)/2$. If $s = 0$, there are no interacting pairs and individuals do not know the phenotypes of any other group members. If $s = 1$, all individuals interact with all other individuals in the group and know whether they are helpers or reproductives. In between these two extremes, individuals may only interact with a subset of the cells in the group and know only partial information about the proportion of helpers in the group.

#### F.3 Individual specialisation

Before we can describe how individuals within the group specialise in this framework, we need to establish a few definitions. For a given interaction network, $G$, let $V = \{v_1, \ldots, v_n\}$ be the set of nodes (individuals or cells) and $E$ be the set of edges, such that $e_{ij} \in E$ if individuals $v_i$ and $v_j$ interact. We define an *allocation of labour* as the map $a : V \to \{0, 1\}$ such that $a(v_i) = 1$ if individual $v_i$ is a helper and $a(v_i) = 0$ if $v_i$ is a reproductive. The realised proportion of helpers to reproductives in the group is then given by $p = \sum_{i=1}^{n} a(v_i)/n$. We denote by $N_G(v_i)$ as the open neighbourhood of individual $v_i$. The degree of $v_i$, $d_i$, is the number of cells in its open neighborhood, that is, $d_i = |N_G(v_i)|$. Using these definitions we can write the *open proportion of helpers in the neighbourhood* of individual $v_i$ as $p_{(i)} = \sum_{v_j \in N(v_i)} a(v_j)/d_i$, that is, the fraction of its neighbours that are helpers. If an individual has no neighbours ($d_i = 0$), then the open proportion of helpers is not defined and the allocation of labour proceeds differently.

For a particular group (given a set of cells $V$ and an interaction network $E$), we determine the allocation of labour, $a$, using a threshold model. First, we assume that the group begins with an initial allocation of all reproductives ($a(v) = 0, \forall v \in V$). This implicitly presupposes that reproduction is the default phenotype of cells. Then for $\tau \geq n$ time steps, we randomly sample with replacement one cell $v_i \in V$ at a time from the group. We sample with replacement and set $\tau \geq n$ so that a cell that has "committed" to a particular phenotype may still switch if too many of its neighbours have adopted the same choice. This emphasises that the allocation of labour process is dynamic and that developmental trajectories are plastic/responsive to their social environment. Each chosen cell, $v_i$, considers the open proportion of cells in its neighbourhood $p_{(i)}$ and chooses to adopt a helper or reproductive phenotype depending on its target proportion of helpers, $t$. If $p_{(i)} < t$, then the cell adopts a helper phenotype ($a(v_i) = 1$) as there are fewer helpers in its neighbourhood than its target proportion of helpers. If $p_{(i)} > t$, then the cell adopts a reproductive phenotype ($a(v_i) = 0$) as there are more helpers in its neighbourhood than its target proportion of helpers. If the open proportion of helpers is equal to the target proportion of helpers ($p_{(i)} = t$), then the cell adopts either phenotype with equal probability. If the individual has no neighbours ($d_i = 0$), then the individual behaves as a random specialist with helper probability equal to the target proportion of helpers ($q = t$).

#### F.4 Fitness calculation

Once labour is allocated within the group, we calculate the fitness of the group using the fecundity equation of the linear public goods model (S7) with $b = 1 - \epsilon$ and $h = \epsilon$, where $\epsilon$ is a measure of how essential cooperation is. We assume that the cost of coordination is given by $c_{\mathrm{C}} = c(1 - e^{-\chi sn(n-1)/2})$, where $sn(n-1)/2$ is the expected number of cell-to-cell interactions, $c \in [0, 1]$ controls the scale of the cost, and $\chi > 0$ determines how diminishing the cost is. We assume that the cost of coordination depends directly on the expected number $sn(n-1)/2$ of cell-to-cell interactions rather than the realised number of cell-to-cell interactions because the former captures the

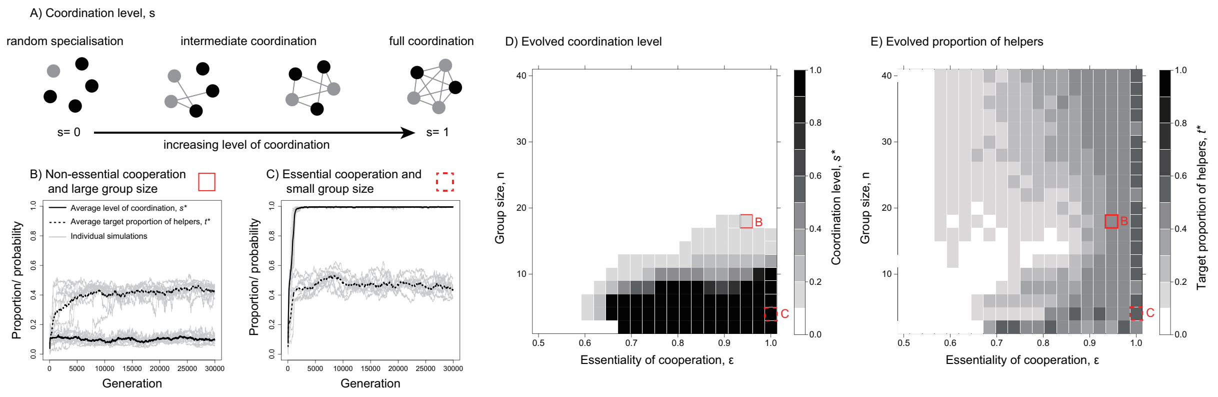

Figure S4: **The optimal level of coordination**. (A) If the level of coordination may evolve, then a spectrum of possible mechanisms could arise. At one extreme ($s = 0$), individuals may not coordinate at all (random specialisation). At the other extreme ($s = 1$), all individuals coordinate with one another and the variance in the realised proportion of helpers is minimised (fully coordinated specialisation). In between ($0 < s < 1$), intermediate forms of coordination may arise in which not all individuals coordinate with one another and sub-optimal proportions of helpers can occur. (B and C) We present the across-simulation average level of coordination ($\bar{s}$, *solid line*) and target proportion of helpers ($\bar{t}$, *dashed line*) that evolves. We also plot individual simulation time series for the trait values (*gray lines*), where each line corresponds to a population average. We find that the traits converge stably within hundreds of generations. Moreover, the evolutionary outcomes (estimated by $s^*$ and $t^*$), depend on demographic and environmental conditions (panel B corresponds to the *red box, solid line* in panel D, whereas panel C corresponds to the *red box, dashed line* in panel D). Separate simulations show that these results are independent of starting conditions. (D) Smaller groups (lower $n$) and more essential cooperation (higher $\epsilon$) favour the evolution of fully coordinated specialisation ($s^* \approx 1$). Intermediate coordination ($0 < s^* < 1$) may evolve in more moderate conditions. (E) Higher essentiality of cooperation (higher $\epsilon$) leads to a higher proportion of helpers. Group size ($n$) has less of an effect on the target proportion of helpers. $c = 0.1$ and $\chi = 0.01$.

effort that each individual puts into coordination instead of how successful its efforts actually were, and so is under evolutionary control.

#### F.5 Creating a new generation

Each generation, we create the new population by sampling group founders from individuals of the previous generation. That is, we randomly pick individuals with a probability proportional to the relative fitness of each group. The process repeats $N_{\mathrm{g}}$ times with replacement. Each sampled individual forms a group of $n$ individuals with inherited trait values $s$ and $t$. When the founder individual is sampled, we assume that there is a chance $\mu$ of having a mutation in one of the traits. If a mutation happens, the trait value is perturbed by adding a normally-distributed random number, $\delta \sim \text{Normal}(0, \sigma^2)$, where $\sigma^2$ is the variance in the size of the mutation, truncated between zero and one.

#### F.6 Simulation results

We perform simulations for each combination of group size $n \in \{2, 4, \ldots, 38, 40\}$ and essentiality of cooperation $\epsilon \in \{1/2, 1/2 + 1/38, \ldots, 1/2 + 18/38, 1\}$. For each simulation we determine the number of groups as $N_{\mathrm{g}} = \lceil N_{\mathrm{T}}/n \rceil$, where $N_{\mathrm{T}} = 10,000$. All individuals are initialised with no coordination or no propensity for division of labour (i.e., $s = t = 0$). Next, we proceed in each generation by determining the interaction network of each group (given its strength of coordination, $s$), and then each group's allocation of labour (given its target proportion of helpers, $t$). We then calculate the fitness of each group and sample individuals for the next generation with a probability proportional to fitness. We assume that mutations occur with probability $\mu = 0.001$ and that the expected variance in the mutation size is $\sigma^2 = 0.1$. We set $c = 0.1$ and $\chi = 0.01$ (parameters of the cost of coordination).

The results of the simulations are shown Fig. S4, where we examined 400 combinations of group size and essentiality and we ran the simulation 10 times for each parameter combination, and where each simulation lasted 30,000 generations. For each simulation, we denote by $\hat{s}$ and $\hat{t}$ the average trait values across the population for a given generation. We denote by $\bar{s}$ and $\bar{t}$ the across-simulation average of $\hat{s}$ and $\hat{t}$ for each generation. Finally, the evolutionary outcomes, $s^*$ and $t^*$, are estimated as the average $\bar{s}$ and $\bar{t}$ in the last 3,000 generations of the simulations. We show the evolutionary outcome $s^*$ as the main results but also plot 10 individual time series of $\hat{s}$ and $\hat{t}$ and the across-simulation average $\bar{s}$ and $\bar{t}$. To account for simulation stochasticity, we categorised random specialisation as any strategy for which $s^* < 0.1$, fully coordinated specialisation as any strategy for which $s^* > 0.9$ and intermediate specialisation as any strategy for which $0.1 < s^* < 0.9$. In Fig. S4, we show the results for $s^*$ and $t^*$ in these simulations, finding that smaller group sizes (lower $n$) and higher essentiality of cooperation (higher $\epsilon$) favours higher levels of coordination (higher $s^*$) and that more essential cooperation (higher $\epsilon$) favours a higher proportion of helpers (higher $t^*$). This is in agreement with the results of the previous analyses, while also highlighting that the level of coordination can also be a factor shaped by natural selection.

### G Dividing labour in a cyanobacteria filament

Here we develop a more detailed model based on the biology of a growing cyanobacteria filament. This model allows for local benefits from a public good in a spatially structured group, and illustrates how local coordination can occur through a signal and response mechanism. While this model is anchored to the particular details of a growing filament, it is also intended as an approximation for more complex spatial distributions.

In the following, we give more details on the cyanobacteria model. First, we specify how filaments increase in cell number as reproductives grow and divide. Second, we detail how the signal-response system can produce coordinated groups. Finally, we describe the evolutionary process used in our simulations to determine the optimal mechanism in different scenarios. The results of this model are depicted in Figs. 4 and 5 of the main text, and in Fig. S6.

#### G.1 Life cycle

Consider a growing filament (a one-dimensional array of connected cells). At time $t \geq 0$, we let $L_t$ be the number of cells in the filament, $I_t = \{1, \ldots, L_t\}$ be the set of (indexes to) individuals in the filament, and $H_t \subset I_t$ and $R_t \subset I_t$ be, respectively, the set of helpers and reproductives. At the start of the filament growth ($t = 0$), we assume that groups consist of four cells, where the two interior cells are reproductives and the exterior cells are helpers. That is, we have $I_0 = \{1, 2, 3, 4\}$, $H_0 = \{1, 4\}$, and $R_0 = \{2, 3\}$.

##### G.1.1 Reproductives absorb fixed $N_2$ to grow

Over time, reproductives increase in size at a rate that depends on the density of the public good that can be absorbed at their location along the filament. We assume the density of the public good at location $i \in R_t$ at time $t$ is equal to

$$\Phi_i^t = \phi + \sum_{j \in H_t} \phi_{i,j}^t, \tag{S44}$$

where $\phi \geq 0$ is the uniform background density of the public good due to the environment, and $\phi_{i,j}^t \geq 0$ is the increase in the local density of the public good that is due to helper $j$ at time $t$, which is assumed to be given by

$$\phi_{i,j}^t = \bar{\phi}(1-s)^{\zeta} \frac{\eta^{|i-j|}}{\sum_{k \in I_t} \eta^{|k-j|}}, \tag{S45}$$

where $\bar{\phi}$ is the maximum rate of public good production by a helper, $(1-s)^{\zeta}$ is the degree to which this production decreases due to a tradeoff with the production of signalling molecules, and the remainder ensures that $\phi_{i,j}^t$ declines by a factor of $0 < \eta \leq 1$ for every cell position that separates the reproductive cell $i$ from the helper $j$ (diffusion factor of fixed $N_2$). When $\eta$ is small, helpers only provide substantial benefits to their nearest neighbours. When $\eta$ is large, even reproductives at a considerable distance along the filament receive benefits. The denominator enforces a conservation principle such that an increase in $\eta$ does not artificially increase the amount of the public good produced by helpers. In the limit as $\eta \to 1$, all reproductives benefit equally from the efforts of each helper.

##### G.1.2 Tracking the size of reproductives

Let $\pi_i^t$ be the size of reproductive cell $i \in R_t$ at time $t$. We assume that the reproductives at the start of group growth cycle start with base size $\pi_i^0 = 0$. For any time interval $\Delta t$ during which no cell divides anywhere in the filament, the increase in size of a reproductive cell $i \in R_t$ is calculated simply as

$$\pi_i^{t+\Delta t} = \pi_i^t + \Psi_i^t \Delta t,$$

where $\Psi_i^t$ is the instantaneous growth-rate of reproductive $i \in R_t$. We assume that $\Psi_i^t$ is an increasing but diminishing function of the rate of public good that is absorbed at its location, $\Phi_i^t$, according to the functional form

$$\Psi_i^t = \psi(1 - e^{\mu \Phi_i^t}), \tag{S46}$$

where $\psi$ is the maximum growth-rate, and larger $\mu$ leads to a more diminishing curve.

##### G.1.3 Replication of reproductives

Reproductives grow until they reach a critical size $\bar{\pi}$, at which point they divide by budding off a daughter cell to one side of the parent cell along the filament. At any time, $t$, we calculate the time until the next replication, $\tau^t$, with a simple procedure. For each reproductive cell $i \in R_t$, we calculate its expected time until replication as

$$\tau_i^t = \frac{\bar{\pi} - \pi_i^t}{\Psi_i^t}, \tag{S47}$$

which is simply the amount it has left to grow divided by its growth rate and where we have held fixed the growth/replication of all other reproductives. Thus, the next reproductive to divide is simply the reproductive with the smallest expected time to replication, $\tau^t = \min_{i \in R_t} \tau_i^t$.

When a reproductive cell $i \in R_t$ divides, it buds off a daughter cell either before or after the parent cell along the filament with equal probability. The positions, phenotypes and sizes of all cells of all other cells in the filament are reindexed to account for the new cell (all cells to the right of the daughter cell move one space further along the indexing array). Whether the daughter cell becomes a a helper or a reproductive depends on the mechanism of specialisation (see section G.2). The parent cell size is set to zero and, if the daughter is a reproductive, its size is set to zero as well. The group life cycle ends when the filament has reached a group size of $L$ cells, at which point all reproductives produce a great number of offspring that disperse to found filaments in the next generation of the group life cycle. The remaining cells then die (non-overlapping generations).

#### G.2 How cells specialise

When a new cell is produced, whether the cell becomes a helper or a reproductive depends on the mechanism of specialisation. There are four co-evolving traits in our model that combined determine the mechanism of specialisation: (i) the baseline probability, $0 \leq q \leq 1$, (ii) the level of signalling, $0 \leq s \leq 1$, (iii) the response sensitivity, $v \geq 0$, and (iv) the response threshold, $d \geq 0$. The baseline probability, $q$, is the probability that the new cell adopts a helper role in the absence of coordination (i.e., when either $s = 0$ or $v = 0$). If the signalling level is positive (i.e., $s > 0$), then helper cells produce signalling molecules that diffuse along the length of the filament. Let $i$ be the position of the new cell at time $t$ in the filament and where the indices of all other cells have been updated to account for the new cell. The level of the signal detected by the cell is

$$\chi_i^t = \max\left(\lambda s \sum_{j \in H_t} \xi^{|i-j|} + \varepsilon, 0\right), \tag{S48}$$

where $\lambda s$ is the maximum rate of signal produced by each helper, and $\xi$ is the factor by which the signal declines for each position that separates the helpers from the receiver. The $\varepsilon$ term is a normal random variable with mean 0 and variance $\sigma_\epsilon^2$ that accounts for the fact that new cells do not perfectly detect the level of the signal.

The degree to which the new cell responds to the signal when specialising depends on an interaction between the detected level of the signal, $\chi_i^t$ and the trait values $q$, $d$, and $v$. Specifically, we model the probability, $p$, that the cell adopts a helper phenotype via the response norm

$$p = \min\left(1, \max\left(0, q + 1 - \frac{2}{1 + e^{-v(\chi_i^t - d)}}\right)\right). \tag{S49}$$

The form of this function is depicted in Fig. 4B in the main text. We assume that if helpers emit no signal, then the response threshold $d$ is also held at zero (i.e., $d = 0$ if $s = 0$). Random specialisation occurs if cells are insensitive to the signal (i.e., $v = 0$) or if cells send no signal at all (i.e., $s = 0$). In this case, equation (S49) becomes $p = q$, and new cells adopt a helper phenotype with the baseline probability $q$. Coordination occurs when $v > 0$ and $s > 0$, in which case the probability of adopting a helper phenotype is affected by the detected level of the signal, $\chi_i^t$. The larger the response sensitivity, $v$, and the larger the difference between the response threshold, $d$, and the detected level of the signal, $\chi_i^t$, the more the probability of adopting a helper phenotype $p$ is perturbed from the baseline probability $q$. If the detected signal level is greater than the response threshold (i.e., if $\chi_i^t > d$ holds), then sensitive cells (with $v > 0$) decrease their probability of adopting a helper phenotype (and thus $p < q$). If the detected signal level is less than the response threshold (i.e., if $\chi_i^t < d$ holds), then sensitive cells (with $v > 0$) increase their probability of adopting a helper phenotype (and thus $p > q$). In the limit as cells become infinitely sensitive ($v \to \infty$), the response norm leads to a deterministic response where if the detected signal level is smaller than the threshold, the new cell always becomes a helper ($p = 1$), and if the detected signal level is greater than the threshold, the new cell always becomes a reproductive ($p = 0$).

In our model, higher coordination incurs larger metabolic costs. First, the more that helpers produce the signal (higher $s$) the less resources they have to produce the public good (see equation (S45)). Second, we assume that filaments with more sensitive cells (higher $v$) grow more slowly, which we model as a an increase in the growth-cap of reproductives, via

$$\bar{\pi} = \bar{\pi}_0 + e^{\beta v} - 1, \tag{S50}$$

where $\bar{\pi}_0$ is the baseline growth-cap. More sensitive cells (higher $v$) lead to an exponentially increasing growth-cap with shape parameter $\beta$.

Consequently, in different scenarios, the optimal trait values $q$, $s$, $d$ and $v$, will depend on the tradeoffs as the filament grows between producing more helpers and producing more reproductives, and between the growth costs and the advantages of coordination.

#### G.3 Evolving filaments

For a given set of model parameters, we determine the optimal trait values $q$, $s$, $d$ and $v$ by simulating an evolving population of cyanobacteria filaments over 4,000 generations. We initialise the population with the resident trait values all equal to zero ($q = s = d = v = 0$). Each generation, we consider an invading mutant strategy, which we draw from a multivariable random distribution $\mathcal{N}(\boldsymbol{\mu}, \boldsymbol{\Sigma})$ with location $\boldsymbol{\mu}$ equal to a vector containing the resident trait values $(q, s, d, v)^\top$ and with covariance matrix:

$$\boldsymbol{\Sigma} = \begin{pmatrix} \sigma_q^2 & 0 & 0 & 0 \\ 0 & \sigma_s^2 & 0 & 0 \\ 0 & 0 & \sigma_d^2 & 0 \\ 0 & 0 & 0 & \sigma_v^2 \end{pmatrix}. \tag{S51}$$

We do not allow all traits to mutate at the same time. For the first 500 generations, we consider only mutations in the baseline helper probability $q$, by setting $\sigma_s^2 = \sigma_d^2 = \sigma_v^2 = 0$. This allows the population to evolve to the optimal strategy for random specialisation. For the rest of the generations, we allow only the coordination traits $s$, $d$ and $v$ to mutate by removing the constraint on $\sigma_s^2$, $\sigma_d^2$ and $\sigma_v^2$ and by instead constraining $\sigma_q = 0$. We set the resident trait value of $q$ to be its average trait value over generations 250 to 500. In addition, we set $d = 0$ whenever $s = 0$.

For each generation of the simulation, we estimate the average fitness of the resident strategy and the mutant strategy by simulating 200 filaments using each strategy. For a given simulation, let $\tau_L$ be the time at which the $L$-th cell in the filament is produced. We calculate fitness as:

$$w_{\text{filament}} = \frac{\sum_{i \in R_{\tau_L}} \Psi_i^{\tau_L}}{\tau_L}, \tag{S52}$$

that is, as the sum of the fecundities of the reproductives in the last generation of the group life cycle (estimated as their growth rates), divided by the time it took the filament to grow to that size. If the average fitness of the mutant strategy is greater than the average fitness of the resident strategy, we replace the resident strategy with the mutant strategy before proceeding to the next generation. If the average fitness of the mutant strategy is less than the average fitness of the resident strategy, we keep the resident strategy when proceeding to the next generation. We estimate the evolved trait values of each evolutionary simulation as the average of each trait value over the last 2,000 generations of the simulation.

#### G.4 Simulation main results

There are 15 parameters in our model (see Table S1). We focused our investigation to the particular patterns produced by two of these parameters, namely $\phi$ and $\eta$. The other parameters and simulation parameters were fixed

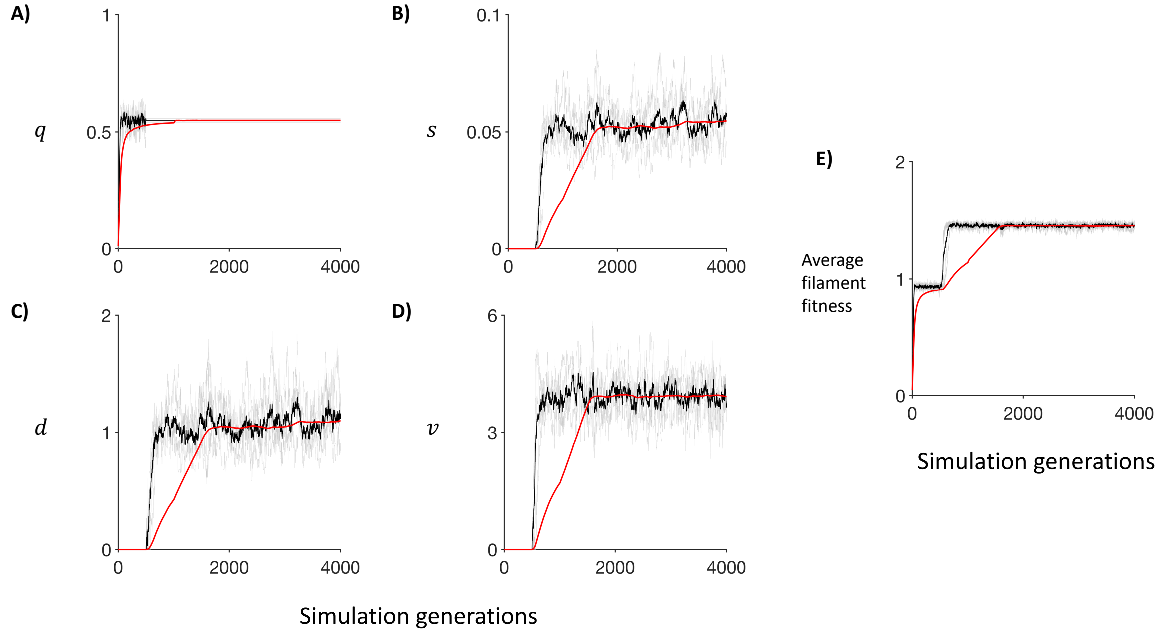

Figure S5: **Convergence to optimal trait values**. We show the convergence of trait values and average filament fitness across 5 independent simulations for the case of essential cooperation ($\phi = 0$) and very local cooperation ($\eta = 0.1$). A) Convergence of the helper probability, $q$. B) Convergence of the signal production, $s$. C) Convergence of the signal threshold, $d$. D) Convergence of the signal response sensitivity, $v$. E) Convergence of average filament fitness. In each case, the grey lines show the trait and fitness values for each of the 5 independent simulations, the black line shows the average trait value or fitness across the 5 independent simulations for that particular generation, and the red line shows the rolling average over the last 1000 generations (truncated if simulation generation is less than 1000). Combined, these show that the rolling averages are stable by the end of the simulations, and that the trait values have approximately converged to the final rolling average within 100-200 generations of being allowed to evolve. The convergence of average filament fitness is particularly stable.

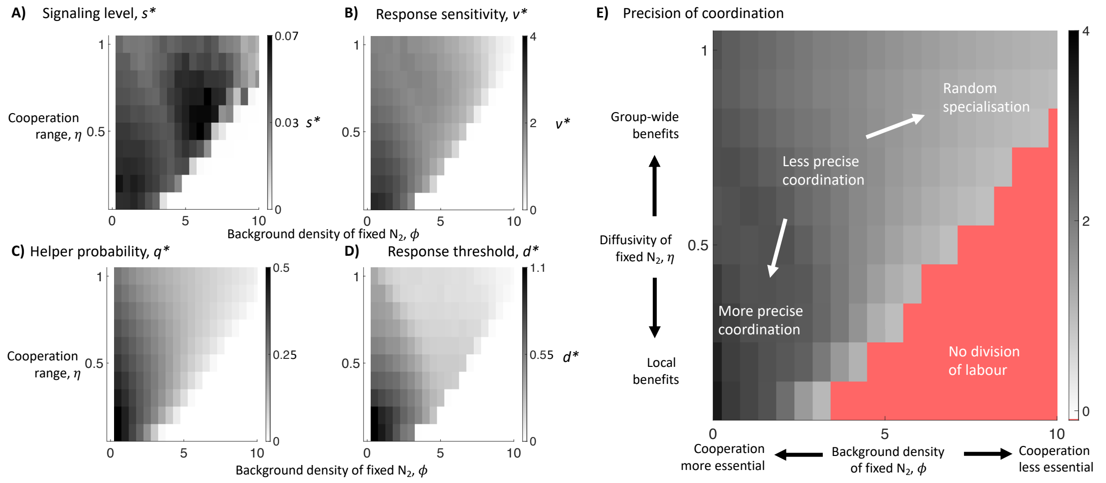

Figure S6: **Simulation results for alternate starting conditions**. We repeated the main simulations analysis while assuming that filament spores consist of 4 cells with no helpers (i.e. a sequence of 4 reproductives: R-R-R-R). We found the same qualitative results as for the main analysis (Fig. 5 in the main text): a lower background density of fixed $N_2$ (smaller $\phi$) and more limited diffusion of fixed $N_2$ (smaller $\eta$) produced filaments that evolved to a relatively more precise allocation of labour (higher coordination).

to values given in Table S1. In Fig. S5, we provide some sample plots to illustrate the evolutionary convergence of our simulations for the specific case of essential cooperation ($\phi = 0$) and very local cooperation ($\eta = 0.1$). We see that all trait values converge to the approximate final rolling average within 100-200 generations of being allowed to evolve. The results for the evolved level of signalling, $s^*$, response sensitivity, $v^*$, baseline helper probability, $q^*$, and response threshold, $d^*$ are shown in Fig. 5 in the main text. In these results, the evolved trait values ($q^*, s^*, d^*$ and $v^*$) are averages across 5 independent evolutionary simulations for each parameter combination, where each individual simulation lasted 4000 generations and were the evolved trait values were averaged over the last 2000 generations.

We performed further simulations to estimate the variance in the allocation of help, that is, we asked to what extent coordination leads to a more precise division of labour. For each parameter combination examined in the main text, we ran 10,000 independent simulations of cyanobacteria growth using the evolved strategies ($q^*, s^*, d^*$ and $v^*$; Fig. 5A-5D in the main text) In each simulation, we recorded both the total number of helpers in the filament and the number of helpers in the leftmost non-terminal (excluding the outside helper) 10 cells in the last generation of each simulated filament. Using this data, we calculated the variance in the number of helpers in the leftmost non-terminal 10 cells of the filament across all simulations with the same parameter combination. We calculated the relative variance as this variance divided by the variance that would be expected of a binomial distribution with 10 trials and probability of success equal to the average proportion of helpers across the 10,000 independent simulations. We approximated the relative precision of a given strategy as the reciprocal of the estimated relative variance. In this way we were able to estimate the degree of precision of the allocation of labour in filaments with use of a single metric (Fig. 5E in the main text).

We also considered the possibility that filaments begin with no helpers. We repeated the above analyses, while ignoring the parameter combinations in which cooperation is essential ($\phi = 0$). Results for the evolved trait values and relative precision are given in Fig. S6, showing the same qualitative patterns.

#### G.5 The effect of helper clumping

We ran additional simulation to quantify the propensity and cost of helper clumps. For a given set of parameter values (Table S1), we extracted the evolved trait values from the previous set of simulations and ran $T$ independent simulations of filament growths with the associated evolved strategy ($q^*$, $s^*$, $d^*$, and $v^*$). When examining random specialisation, we set $s^* = d^* = v^* = 0$. Within a single simulation, we define a clump as any contiguous grouping of helpers in the last generation of the group growth. Thus, clump sizes can range from 1 (a single helper) to $L$ (the entire filament). For each simulation, we calculated the average clump size over all clumps in the filament. The average clump size across all $T$ simulations is then the across-simulation average of this within-simulation average. To determine the cost of clumping, we recorded both the fitness of each filament and its average clump size across all $T$ simulations. We then set the "cost of clumping" as equal to the slope of the linear least-squares regression of filament fitness on average clump size.

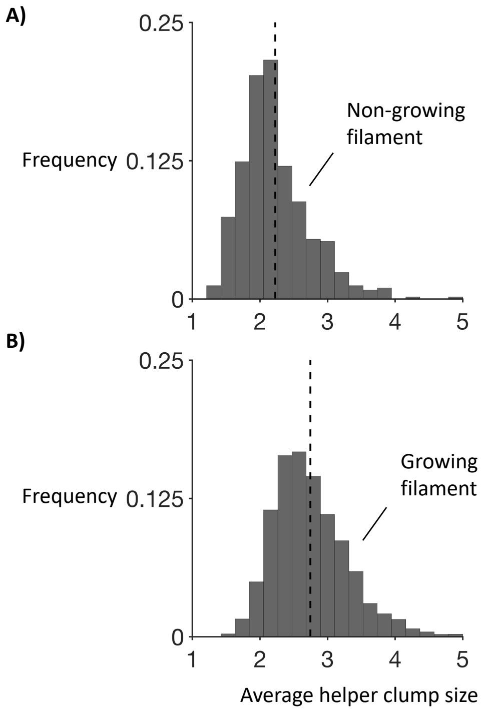

Figure S7: **Growing groups produces larger helper clumps**. We considered the distribution of average helper clump sizes for the cases where: A) helper and reproductive roles are randomly assigned in the final generation of group growth (uncoupling division and differentiation), and B) helper and reproductive roles are assigned as the group grows (division and differentiation coupled). We specifically examined the case where cooperation was essential ($\phi = 0$) and very limited diffusion of fixed $N_2$ ($\eta = 0.1$), performing 10,000 independent simulations for each strategy. We found that helper clumps still occur when division and differentiation are uncoupled but that average clump size increases when differentiation occurs as the group grows (average clump size for uncoupled: $2.23$ and coupled: $2.76$).

In Figure 6A and 6B of the main text, we used the above approach to map the propensity and cost of clumping as a function of the background density of fixed $N_2$ ($\phi$) and the diffusivity of fixed $N_2$ ($\eta$). In this case, we calculated the cost of clumping using relative rather than absolute fitness, where the fitness of the filament in each simulation is divided by the average fitness of all filaments with the same associated parameter values. In Figure 6C and 6D of the main text, we used the above approach to determine the differences between random specialisation and coordinated specialisation, focusing on the extreme case of essential cooperation $\phi = 0$ and very low diffusivity of fixed $N_2$ ($\eta = 0.1$). In this case, the cost of clumping was calculated with the absolute fitness of each filament, as the underlying parameters are the same for both the random and coordinated treatments.

We then sought to determine the effect of group growth on the formation of helper clumps in randomly specialising filaments. To do this, we ran $T$ independent simulations, where filament does not grow and is composed $L$ cells. Individual cells become a helper with a random probability equal to the optimal strategy for the growing group $q^*$ and otherwise become reproductives. We then quantified average clump size in each simulation in the same way as for the previous analysis. Figure S7, shows the distribution of average clump sizes across $T = 10,000$ simulations of non-growing filaments (A) and growing filaments (B). This was calculated for the extreme case of essential cooperation $\phi = 0$ and very low diffusivity of fixed $N_2$ ($\eta = 0.1$). We find that (A) non-growing groups still form helper clumps but that the distribution has a smaller average value and has a smaller upper tail than for (B) growing filaments. Consequently, helper clumping is more severe when cellular division and differentiation are "coupled".

Table S1: **Cyanobacteria model**. Evolutionary traits, model parameters and simulation parameters of the cyanobacteria model, with their definitions and associated values in the presented simulations. Parameters of the evolutionary simulation such as the number of generations per simulation, the mutation rate or size, or the number of independent replicates are not shown here (but see supplementary section G).

| **Evolutionary traits** | | |
|---|---|---|
| **Notation** | **Definition** | **Value(s)** |
| $q$ | Helper probability in the absence of coordination. | $0 \leq q \leq 1$ |
| $s$ | Relative amount of signal produced by helpers. | $0 \leq s \leq 1$ |
| $d$ | Response threshold to signal by new cells. | $d \geq 0$ |
| $v$ | Response sensitivity to signal by new cells. | $v \geq 0$ |

| **Model parameters** | | |
|---|---|---|
| **Notation** | **Definition** | **Value(s)** |
| $\phi$ | Environmental background density of fixed $N_2$. | $\phi \in \{0, 0.5, \ldots, 9.5\}$ |
| $\eta$ | Relative diffusivity of fixed $N_2$ produced by helpers. Lower values means that benefits are shared more locally. | $\eta \in \{0.1, 0.2, \ldots, 1\}$ |
| $\bar{\phi}$ | Maximum rate of $N_2$ production by helpers. | 50 |
| $\psi$ | Maximum growth-rate of reproductives. | 10 |
| $\mu$ | Shape of reproductive growth-rate as a function of fixed $N_2$ intake. Higher values means more diminishing. | 0.25 |
| $\bar{\pi}$ | Critical size at which reproductives replicate (with no response sensitivity.) | 100 |
| $\beta$ | Shape of increase in critical replication size due to higher response sensitivities. Larger values means a more accelerating cost. | 0.5 |
| $\lambda$ | Maximum signal production rate by helpers. | 30 |
| $\zeta$ | Shape of trade-off between signal production and $N_2$ fixation. | 2.5 |
| $\xi$ | Diffusion range of the signal. Lower values means that the signal decays rapidly along the filament. | 0.5 |
| $\sigma_\epsilon^2$ | Noise in local signal detection by new cells. | 0.01 |
| $L$ | Size of filament at end of group life-cycle. | 50 |

| **Simulation parameters** | | |
|---|---|---|
| **Notation** | **Definition** | **Values** |
| $\sigma_q^2$ | Mutation variance in helper probability, $q$ | $1 \times 10^{-3}$ |
| $\sigma_s^2$ | Mutation variance in signal production, $s$ | $1 \times 10^{-5}$ |
| $\sigma_d^2$ | Mutation variance in signal response threshold, $d$ | $1 \times 10^{-2}$ |
| $\sigma_v^2$ | Mutation variance in signal response sensitivity , $v$ | $5 \times 10^{-2}$ |
